# A Human Tumor Virus Recruits HSP70 Chaperone to Remodel Its Replisome on DNA

**DOI:** 10.64898/2026.11.05.764654

**Authors:** Li Wan, Xiaochen Li, Paul Vittot, Jeffrey L. Brodsky, Peter Wipf, Patrick S. Moore, Yuan Chang

## Abstract

Human Merkel cell polyomavirus (MCV) large T (LT) protein functions as a replication helicase analogous to the cellular MCM complex but requires a second MCV protein, small T (sT), for efficient LT-initiated replication. Both LT and sT encode highly-conserved DnaJ domains that bind to the cellular refolding molecule, heat shock protein 70 (HSP70). Using single-molecule imaging and FRET, we find that sT recruits dimeric HSP70 to serve as a molecular bridge between sT and LT on DNA and that this complex facilitates remodeling LT into a functional double-hexameric helicase. Disruption of LT-HSP70-HSP70-sT interaction with HSP70 inhibitors or by mutations to the LT-sT DnaJ domains collapse the LT-HSP70-HSP70-sT complex, abolishing sT-enhanced LT dodecamerization and ATP-dependent DNA unwinding. Assembly of virus-free mammalian MCM3, MCM4 and MCM6 in nuclear extracts similarly recruits HSP70 to DNA suggesting that Hsp70 remodeling is used for efficient eukaryotic assembly of the multi-protein replication machinery.

## INTRODUCTION

DNA replication in both prokaryotic and eukaryotic systems relies on tightly regulated protein-DNA interactions. Early studies demonstrated that λ bacteriophage DNA replication in Escherichia coli requires bacterial heat-shock chaperone proteins, such as DnaK and DnaJ, to remodel the replisome on the phage origin of replication^1–4^.

Polyomaviruses, including the oncogenic human Merkel cell polyomavirus (MCV) causing Merkel cell carcinoma^5^, serve as valuable models for studying eukaryotic DNA replication and, like lambda phage, initiate viral replication at a single origin sequence^6,7^. The polyomavirus large T antigen (LT) initiates replication by assembling as a head-to-head, double-hexameric, homopolymeric helicase complex on viral origin DNA similar to the eukaryotic MCM2-7 heteropolymeric helicase complex (**Fig. 1A**)^8–10^. Once LT is assembled on the viral origin as a double-hexameric helicase, it initiates ATP-dependent DNA unwinding that allows recruitment of cellular replication proteins to duplicate the viral DNA.

**Fig. 1.**
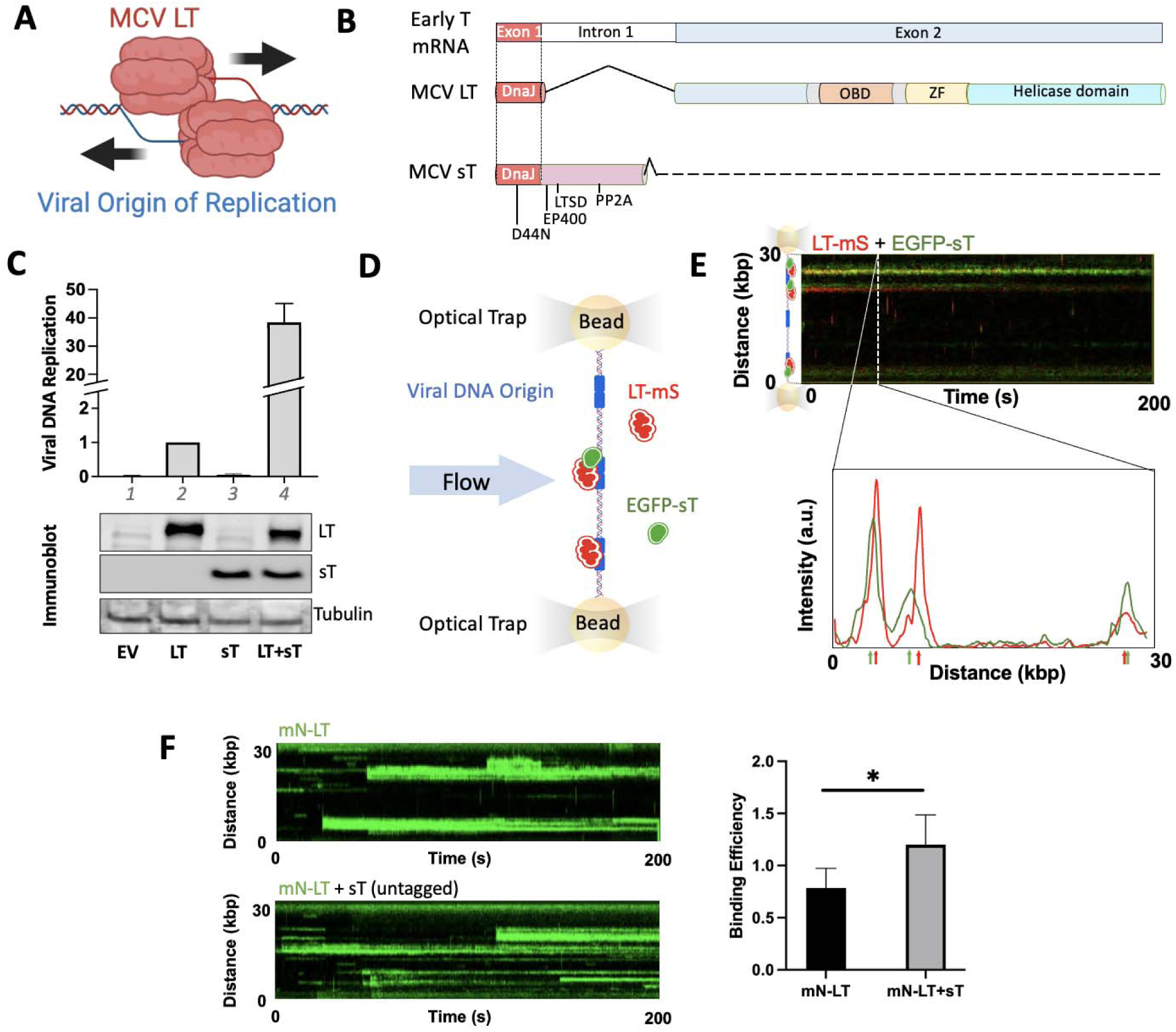
MCV sT promotes LT binding and multimerization on Ori98 DNA. **(A)** Structure of the MCV LT replication head-to-head double-hexamer helicase assembled on the replication origin. **(B)** Functional domains of MCV LT and sT proteins. LT and sT share an N-terminal coding region encoding identical DnaJ domains, disrupted in both by a D44N mutation. LT domains include the origin binding (OBD), zinc-finger (ZF), and helicase domains. sT uniquely encodes EP400- and PP2A-binding motifs and an LT stabilization domain (LTSD). **(C)** sT promotes MCV DNA replication. MCV origin replicon was expressed in all lanes with different combinations of LT and sT in U2OS cells. Upper panel, replicon replication efficiency for LT was increased 38-fold by sT and LT co-expression compared to LT alone. Lower panel, immunoblot confirmation of viral protein expression. Lane 1, empty vector control (EV); lane 2, LT alone; lane 3, sT alone; lane 4, LT+sT. Average and standard deviation for three independent replicates. **(D)** Single molecule DNA binding. C-Trap experimental schematic is shown in which nuclear extracts expressing fluorescent LT-mScarlet (LT-mS, red) and EGFP-sT (green) were mixed in the C-Trap laminar flow channel and flowed against biotinylated multi.Ori98 DNA (blue) tethered between streptavidin beads. Fluorescent protein binding kinetics to origin DNA were determined by confocal microscopy. **(E)** Adjacent LT and sT binding to MCV origin DNA. Representative kymograph is shown in which the fluorescence tracing at a single time point (dotted white line) is cross-wise expanded to reveal adjacent binding between EGFP-sT (green) and LT-mS (red) on the 98 bp viral origin (Ori98). Representative of six independently acquired kymographs. **(F)** Representative kymograph images of mN-LT (green) binding to Multi.Ori98 DNA in the absence (upper) and presence (lower) of untagged sT, and quantification of mN-LT binding efficiency (events·kbp⁻¹·min⁻¹). Mean ± SD, n = 6 kymographs per condition; Two-tailed unpaired Welch’s T test, P=0.0294.

We have previously shown that MCV and SV40 LT initiates DNA initial unwinding through a strand-invasion mechanism that is independent of ATP hydrolysis to form the initial replication bubble.^5^ This mechanism enables the formation of two hexameric LT complexes, one on each origin DNA strand, which then proceed to unwind the DNA in an ATP-dependent manner, facilitating leading and lagging strand synthesis by cellular polymerases^11^. While single molecule studies using nuclear extracts model these early replication bubble formation events, single molecule replisome processivity has not been achieved using T antigens. In cellular assays, however, MCV LT alone has been shown to promote low levels of MCV DNA replication that can be markedly enhanced by a second viral accessory protein called small T (sT) antigen^12^. Small T inhibits Fbw7 that targets LT for proteasome-dependent degradation and in some cell lines increases LT concentration^13,14^. This mass-action effect, however, does not fully account for the potent replication enhancement observed with sT^15^. Despite MCV sT’s known signaling activities and its localization to viral replication foci in infected cells^16–18^, a direct role for sT at the replisome has not been established.

Like other polyomaviruses^19,20^, MCV LT and sT are expressed from alternatively-spliced early transcripts that encode a shared N-terminal region (**Fig. 1B**)^6,21^. This N-terminal sequence encodes a DnaJ domain^22^ that allows both LT and sT to bind to the cellular HSP70 protein that functions as a central hub of the cellular chaperone network^6,19,23–26^. While bacterial DnaK and DnaJ chaperone proteins are known to remodel the bacteriophage DNA replication complex^27^, a comparable role for HSP70 in remodeling the assembled eukaryotic DNA replication machinery is not known. The DnaJ domain of SV40 large T antigen has been shown to promote efficient SV40 DNA replication and has been proposed to facilitate HSP70-dependent rearrangement of LT complexes on viral DNA^28^. However, the precise molecular mechanism underlying polyomavirus LT–HSP70 interaction remains unclear.

Here, we show that presence of MCV sT together with LT promotes MCV DNA replication 38-fold over LT alone. Using C-Trap single-molecule imaging of tandem MCV DNA origins (Multi.Ori98) tethered to polystyrene beads^5^, we show that MCV LT and sT recruit HSP70 via their shared DnaJ domains to origin DNA to enable HSP70-mediated remodeling of the replication complex into a mature, replication-competent LT double-hexamer. Dimeric HSP70 forms a bridge between the LT and sT DnaJ domains on DNA and this HSP70-containing structure remodels LT into dodecamer that efficiently initiates replication. Single-molecule Förster resonance energy transfer (FRET) reveals that MCV LT and sT in this complex are separated by HSP70 dimers beyond the Förster radius, but collapse into direct proximity when either DnaJ domain is mutated or when the complex is treated with HSP70 inhibitors. These data reveal that MCV LT alone is capable of inefficient MCV DNA replication but sT-mediated HSP70 recruitment to DNA-bound LT is required for remodeling LT into an efficient replication helicase. Expanding these findings to cellular MCM helicase proteins reveals that HSP70 is also recruited to DNA suggesting that on DNA remodeling by HSP70 may be a common feature for the eukaryotic DNA replication machinery.

## RESULTS

MCV sT protein enhances LT replisome binding on DNAWhile LT alone inefficiently multimerizes on the viral origin to initiate replication^6,14^, co-expression of LT with sT in U2OS cells increased replication activity up to 38-fold (**Fig. 1C**). To examine whether this enhancement arises from direct association between sT and LT, we performed single-molecule analysis of DNA nuclear extract (SMADNE) assays^29^. When expressed alone, EGFP-tagged sT exhibited nonspecific DNA binding (**Fig. S1A-C, S2**), with a significantly lower kon and lifetime compared to LT dodecameric helicase (**Fig. S3**). However, in the presence of mScarlet (mS)-tagged LT, sT specifically relocalized to the 98 bp MCV origin, forming part of the LT replication complex **(Figs. 1D-E, S1D-F),** resulting in an enhancement of overall observed LT binding efficiency (Fig. 1F). Notably, fluorescence signal tracing revealed a consistent spatial separation between LT and sT binding peaks on the origin, with an average interpeak distance of 196 nm (range: 91-301 nm), corresponding to approximately ∼575 bp of DNA (range: 265-885 bp) (**Fig. S4**); given the diffraction-limited resolution of C-Trap fluorescent imaging (∼250bp), these measurements reflect the time-averaged positions of each labeled protein within its binding zone on DNA rather than precise molecular contact distances.

### sT Enhances LT Binding Affinity and Promotes Replication Competence

Statistics on single-molecular sT-LT interaction revealed the functions of sT in enhancing LT binding. Specifically, co-expression of sT enhanced LT’s binding affinity and assembly on origin DNA (**Figs. 2A, 2B)** with the apparent association rate constant^30^ for mN-LT increasing by 50-75% in the presence of sT (**Fig. 2A**; see **Methods** for kinetic modeling). Notably, sT selectively increased the frequency of replication-competent, dodecameric (12-mer) mN-LT^5^ bound to the origin while generally reducing the frequency of lower-order, non-replication competent LT assemblies (**Fig. 2B**). DNA binding efficiency for mN-LT increased 53% when sT was introduced together with mN-LT, compared to mN-LT alone, but this effect was abolished by mutation of the sT DnaJ domain (sT^D44N^), which disrupts HSP70 binding^6,28,31,32^ (**Fig. 2C, S5**). In contrast, mutations to other sT domains that have been previously found to be responsible for LT-stabilization^14^, PP2A interaction^15^, or EP400-binding^16^ had little or only a marginal effect on LT-DNA binding. Co-immunoprecipitation of bulk LT, sT and HSP70 from U2OS cell extracts further confirmed that these proteins interact in vivo (**Fig. S6**).

**Fig. 2.**
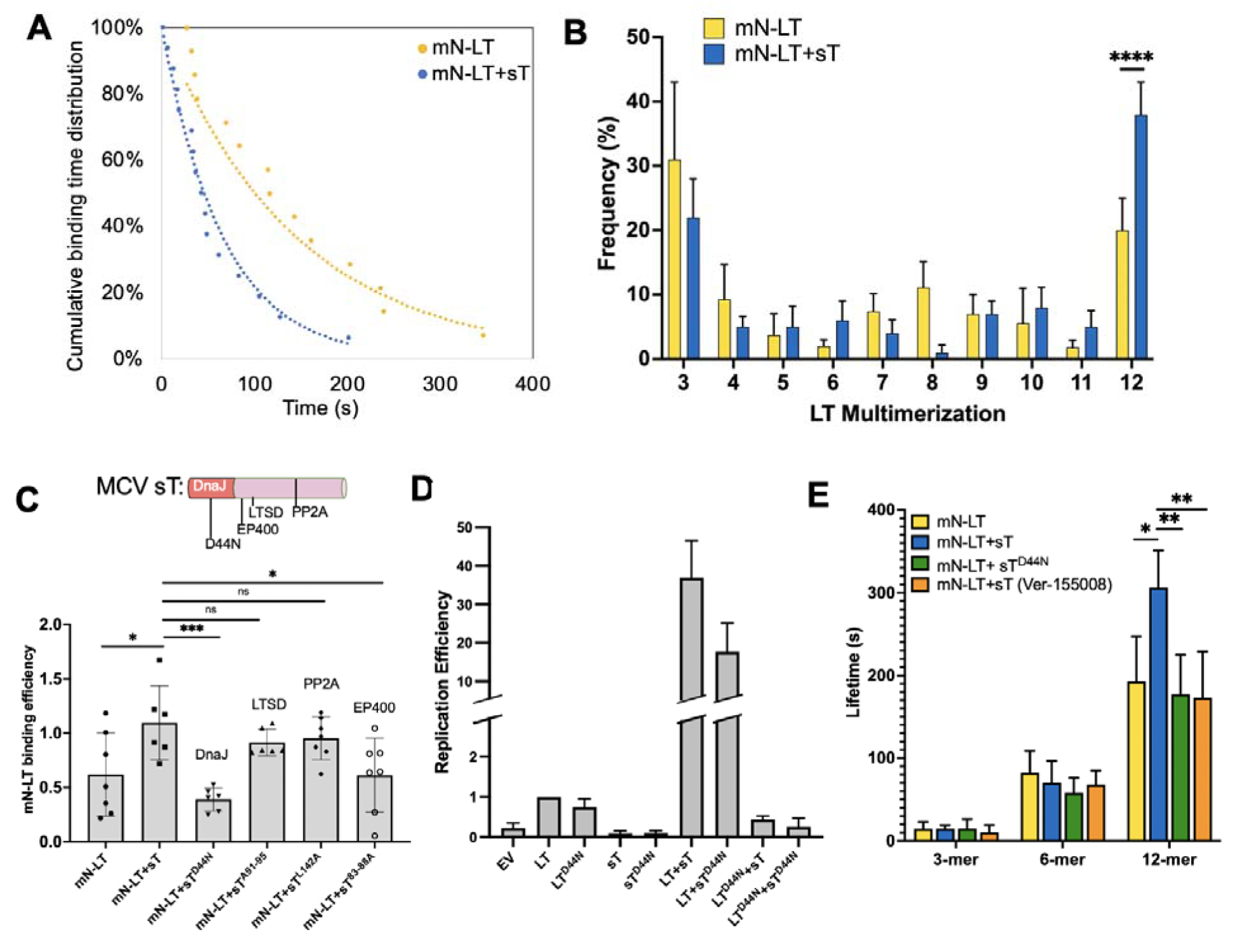
MCV sT DnaJ domain is required for enhanced LT DNA-binding and selectively increases dodecameric LT assembly. **(A)** Small T increases k_on_ for LT binding to viral origin DNA. Cumulative molecule binding time distributions for mN-LT in the presence (blue) or absence of sT (yellow) are shown. Each point represents a single binding event which was fitted to an exponential decay curve to determine kon. In the presence of sT, the estimated LT binding rate increased by 2.40-fold (95% CI, 1.15–4.95; likelihood-ratio test, P = 0.0173). These results suggest that sT promotes LT association with DNA. **(B)** Small T increases LT dodecamerization on origin DNA. Quantitation of LT molecules bound to origin DNA was determined using mNeon-LT (mN-LT) nuclear extracts mixed in situ with untagged sT nuclear extracts. The distribution of mN-LT oligomers bound to Multi-Ori98 DNA in the presence (blue bars) and absence (yellow bars) of sT is shown. Small T selectively enhanced dodecameric (12-mer) LT formation on DNA nearly 2-fold. Two-tail unpaired Welch’s T test for 12-mer shows p<0.0001, (N=6 kymographs per condition.) **(C)** Small T DnaJ mutation eliminates enhanced LT-origin binding. mN-LT origin binding efficiency (binding events•kbp^-1^time^-1^) was enhanced by in situ mixing in the laminar flow channel LT with wild-type sT or LTSD (sT^91–95A^) or PP2A-binding (sT^L142A^) mutants, and only marginally reduced by mixing with the EP400 mutant (sT^83–88A^). Enhancement was lost when sT^D44N^ was mixed with LT. Binding events determined from six kymographs for each condition. **(D)** Optimal MCV origin replication requires HSP70 remodeling of LT. Enhanced LT replication of replicon plasmid in U2OS cells (as in Fig 1B) was markedly reduced using sT^D44N^ co-expression and absent for the LT^D44N^ mutant with or without sT co-expression. Average and standard deviation for three independent replicates. **(E)** Enhanced LT dodecamer binding to origin DNA is lost with sT DnaJ domain mutation or HSP70 inhibition. In situ mixing of sT and LT (blue) increased LT binding lifetime for LT dodecamers but not trimers or hexamers on Ori98 DNA compared to LT alone (yellow). Substitution with sT^D44N^ (green) or pre-treatment with HSP70 inhibitor (5 µM Ver-155008 for 10 min, room temperature, orange) reduced bound dodecamer lifetime to levels seen without sT mixing. Binding events determined from six kymographs for each condition.

Consistent with these findings, bulk viral DNA replication assays in U2OS cells showed that sT^D44N^ reduced replication activity by 52% (**Fig. 2D**) compared to wild-type sT. The LT^D44N^ mutant essentially abolished replication, reducing replication by 99%, even in the presence of wild-type sT, underscoring the critical role of LT-HSP70 interactions in replication enhancement (**Figs. 2D, S7**). Importantly, sT increased the lifetime (τ = 1/K_off_) of the fully-assembled dodecameric mN-LT complex on DNA, without affecting the stability of lower-order LT assemblies, such as trimers and hexamers (**Fig. 2E**). This effect was abolished by the sT^D44N^ mutation or treatment with Ver-155008^33^, a selective HSP70 ATP binding inhibitor (**Fig. 2E**).

### HSP70 Bridges LT and sT on DNA

Given the high cellular concentration of HSP70 typically present in cells^26^, it was likely that non-fluorescent, endogenous HSP70 from C-Trap SMADNE lysate preparations was responsible for mediating the interactions between LT and sT on DNA. To confirm this, we performed three-color fluorescence imaging using EGFP-HSP70 (HSPA1A), HaloTag-JF549-tagged LT, and HaloTag-JF646-tagged sT, all bound to MCV origin DNA (see **Methods** for labelling strategy). These three proteins simultaneously co-localized to origin DNA but were arranged with spatially-distinct and adjacent signals as determined by fluorescence peak tracing (**Fig. 3A, S8**). Within these complexes, the average sT–HSP70 and LT–HSP70 distances were approximately 150 bp and 182 bp, respectively (**Fig. S9**). These sub-complex distances are more consistent with the physical dimensions of an HSP70-bridged assembly, and the larger overall LT–sT separation likely reflects both the diffraction-limited resolution of C-Trap fluorescence imaging and the dynamic positional range of each protein within its binding zone on origin DNA, as has been documented for other DNA-protein complexes studied by single-molecule imaging^34,35^. Photon intensity analysis of HSP70 within DNA-bound HSP70-LT complexes revealed that HSP70 molecule number increases with LT multimerization level, with the greatest HSP70 recruitment observed at LT dodecamers and further enhanced by sT co-expression (**Fig. S10**). Mutations in the DnaJ domains of either LT (LT^D44N^) or sT (sT^D44N^) disrupted the spatial separation between the LT (blue) and sT (red) signals, resulting in a collapse of the characteristic interpeak LT-sT gap (**Figs. 3B, 3C**). These results support a model in which HSP70 dimers bridge LT and sT via their respective DnaJ domains (LT-HSP70-HSP70-sT). While mutant sT and LT molecules still bound together on the origin, suggesting a DnaJ-independent LT-sT interaction surface, proper structural spacing appears to be required for the enhanced LT assembly. We next tested a prediction of this bridging model using Förster resonance energy transfer (FRET). Because FRET reports only whether a donor–acceptor pair lies within the Förster radius (R₀ = 5.9 nm)³⁶, it cannot measure separation across an HSP70 bridge; it provides a threshold test distinguishing the bridged arrangement from a collapsed one. Wild-type mN-tagged LT (donor, excited at 488 nm) and HaloTag-JF549-tagged sT (acceptor) exhibited minimal FRET (<0.5%) when bound to Multi-98 origin DNA, indicating separation beyond the FRET distance limit (R_0_=5.9 nm)^36^ (**Figs. 3D, S11**). In contrast, FRET signals were detected when either LT or sT carried the D44N DnaJ-domain mutation (**Figs. 3E, F, S11**), indicating that disruption of DnaJ-mediated HSP70 engagement alters the spatial organization of the LT–sT complex and brings the labeled fluorophores into closer proximity. Further photobleaching of the acceptor JF549 fluorophore induced dequenching and recovery of the donor mNeongreen signal for D44N mutated sT (**Fig. 3E)** or LT (**Fig. 3F**), but not for wild type (Fig. 3D), confirming that the acceptor signals detected in the mutant conditions arise from donor quenching by energy transfer rather than from spectral bleed-through. These two measurements operate at different scales and are complementary: confocal peak separation (resolution ∼250 bp) positions LT, HSP70 and sT relative to one another within the complex, whereas FRET reports only whether the LT–sT separation has fallen below ∼6 nm. The loss of the resolved LT–sT gap in the D44N mutants (Figs. 3B, 3C) and the appearance of FRET in those same mutants therefore report the same collapse at two different resolutions.

**Fig. 3.**
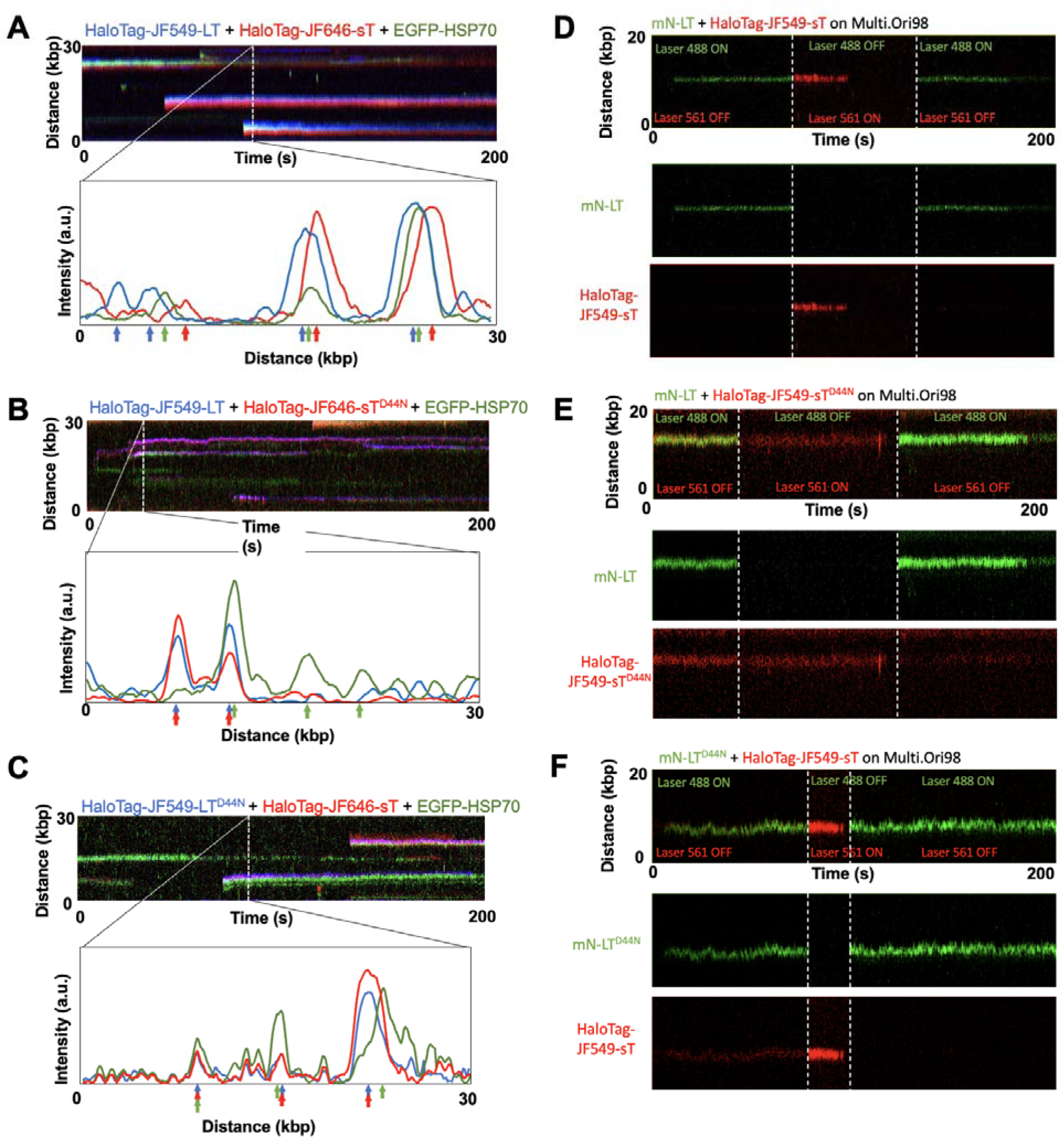
MCV sT forms a scaffold with DNA-bound LT to promote HSP70-mediated LT remodeling into a replication-competent dodecameric helicase. **(A)** HSP70 assembles together with sT and LT on origin DNA. In situ mixing and three-color fluorescence with HaloTag-JF549-LT, HaloTag-JF646-sT and EGFP-HSP70 showed colocalization of all three molecules on origin DNA with peak separations consistent with HSP70 bridging between LT and sT molecules. **(B)** and **(C)** Adjacent molecular assembly of LT, HSP70 and sT to origin DNA was lost with D44N mutation in either sT **(B)** or LT **(C). (D-F)** Forster resonance energy transfer (FRET) analysis of mN-LT and HaloTag-JF549-sT. mN-LT served as the donor and HaloTag-JF549-sT served as the acceptor. DNA-binding events were recorded with 488-nm and 561-nm excitation to detect mNeonGreen and JF549 signals, respectively. After the 561-nm laser was turned off (red dotted line), any remaining JF549-channel signal would indicate FRET from mNeonGreen excitation. The JF549 acceptor was then bleached to completion at 561 nm and donor emission re-measured under 488-nm excitation, where recovery of mNeonGreen signal reports FRET-dependent donor quenching. (**D**) For wild-type mN-LT and HaloTag-JF549-sT, the acceptor-channel signal under donor-only excitation was not detected and acceptor photobleaching produced no donor recovery; this panel therefore shows, within a single trace that the residual acceptor signal is spectral rather than FRET. FRET signal, together with donor recovery on acceptor photobleaching was detected for mN-LT/HaloTag-JF549-sT^D44N^ (**E**) and mN-LT^D44N^/HaloTag-JF549-sT (**F**). (**G**) Kymograph of mNeonGreen bleed-through into the JF549 channel under 488-nm excitiation alone. The corresponding photon counts, which define the background baseline for (D-F), are given in Fig. S10. Display settings as in Methods.

### HSP70 and sT Remodel LT Into a Functional, ATP-Dependent Dodecameric Helicase

Quantitative single-molecule analysis of LT multimerization demonstrated that the presence of sT fundamentally alters the assembly of LT at the viral origin by nearly doubling the formation of functional dodecameric mN-LT helicase complexes compared to mN-LT alone (**Fig. 2B**). Importantly, substitution of wild-type sT with sT^D44N^ mutant reduced dodecamer formation to levels seen with LT alone (**Fig. 4A**), consistent with sT-directed chaperone remodeling of the LT complex. Remodeling activity also required functional HSP70. Pharmacological treatment of nuclear extracts with HSP70 inhibitors, including the ATP-competitive inhibitor Ver-155008^33^, the DnaJ-binding allosteric inhibitor MAL3-101^37^, or the nucleotide exchange inhibitor MKT-077^38^, abolished the sT-mediated enhancement of dodecamer formation (**Fig. 4A**). In contrast, when LT was expressed alone, neither HSP70 inhibition nor the LT^D44N^ mutation had a significant impact on the frequency of dodecamer formation (**Fig. 4A**).

**Fig. 4.**
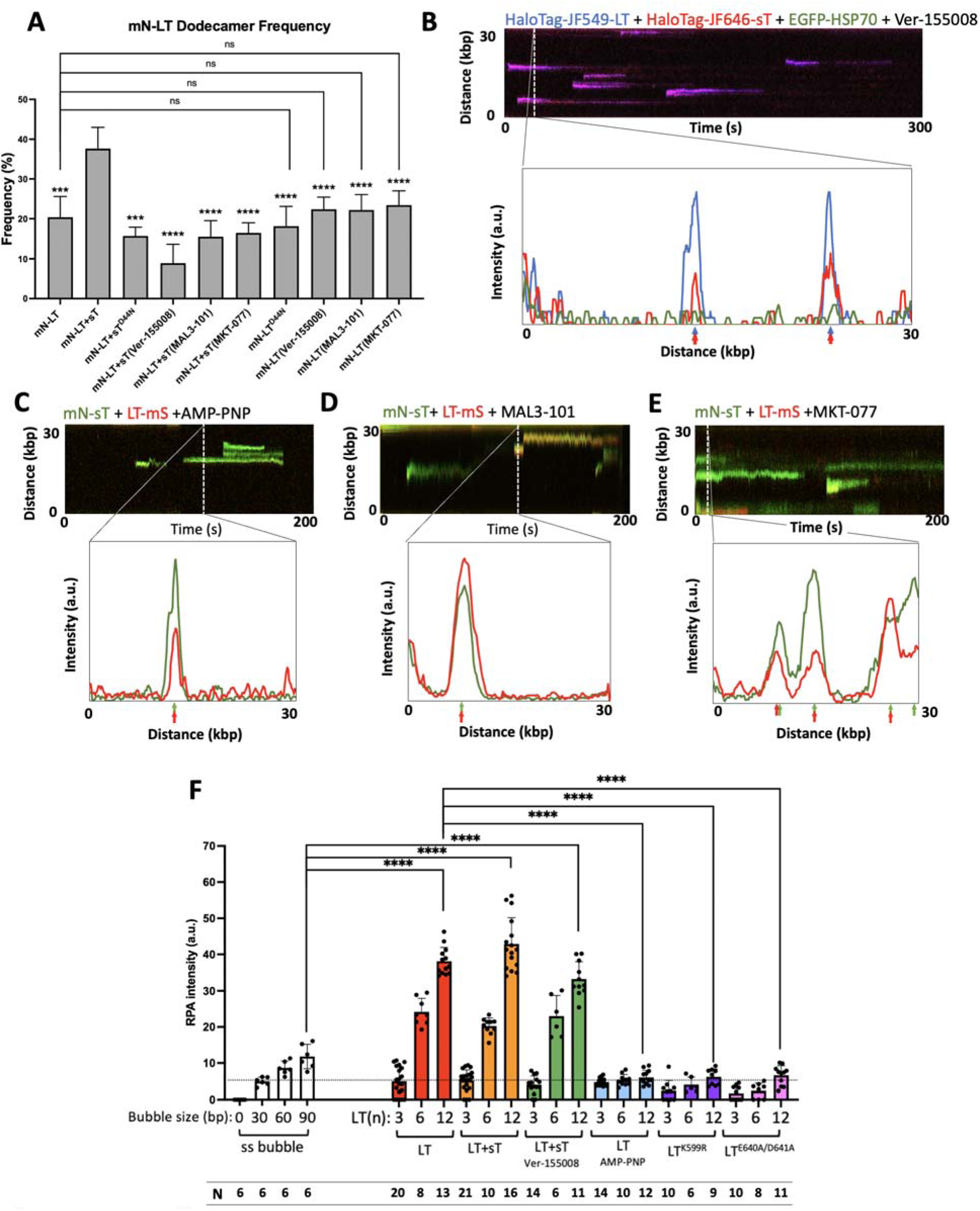
HSP70 inhibition reduces the efficiency of LT assembly on DNA into a replication competent helicase. **(A)** MCV small T promotes LT dodecamerization remodeling through HSP70 interaction. LT dodecamer frequency among total Ori98 DNA binding events was nearly doubled by presence of sT protein. Mutation at the sT^D44N^ or pretreatment with HSP70 inhibitors eliminated the sT-induced LT remodeling. D44N mutation of LT or pre-treatment with Ver-155008 did not significantly reduce dodecamer formation compared to the base level for LT expression alone. Data shown based on total binding events from seven kymographs for each condition. **(B)** HSP70 inhibition prevents HSP70 recruitment to DNA-bound LT-sT and collapses the LT-sT structure. DNA-bound LT still interacted with sT protein (blue+red) but failed to recruit HSP70 (green) to the helicase complex. In contrast to absence of inhibitor (Fig. 2B), HSP70 inhibition was associated with overlapping LT and sT fluorescence. Representative of six kymographs for HaloTag-JF549-LT (blue), and HaloTag-JF646-sT (red) and EGFP-HSP70 (green) after 5 µM Ver-155008 pre-treatment. **(C)** The LT-sT remodeling structure requires active ATP hydrolysis. Representative of six kymographs showing LT-mS (red) and mN-sT (green) binding following apyrase-mediated ATP depletion and addition of non-hydrolyzable ATP analog AMP-PNP (1 mM). AMP-PNP induced collapse of LT and sT binding to DNA. **(D-E)** Other HSP70 active site inhibitors generate similar overlap transitions for DNA-bound LT-sT confirming the requirement of active HSP70 for formation of the LT-sT remodeling structure on DNA. Kymograph showing LT-mS (red) and mN-sT (green) binding to Multi-Ori98 DNA after treatment with HSP70 inhibitors MAL3-101 (50 µM, **D**) and MKT-077 (50 µM, **E**). **(F)** Quantitation of exposed single-stranded DNA (ssDNA) generated by mN-LT 3,6, and 12-mer under the indicated conditions, measured by the fluorescence intensity of HaloTag-JF646-labeled replication protein A (RPA). RPA fluorescence was calibrated using λDNA and λDNA substrates containing defined 30-, 60-, and 90-bp poly(dA) bubbles to estimate the length of exposed ssDNA. Dotted line denotes the RPA intensity at 30-np poly (dA).

At the single-complex level, disruption of HSP70 activity destabilized the origin-bound LT-sT structure. Pre-incubation with Ver-155008 resulted in dissociation of HSP70 from the complex, followed by collapse of both LT and sT signals on DNA (**Fig. 4B**). Complex disruption could also be mimicked by replacing ATP with AMP-PNP, a non-hydrolyzable ATP analogue that supports LT-DNA binding^5^, indicating the necessity of ATP hydrolysis for HSP70 chaperone activity for LT remodeling (**Fig. 4C**). Similarly, pre-treatment with MAL3-101 or MKT-077 resulted in collapse of the LT-sT structure (**Figs. 4D,4E**).

To demonstrate that the dodecameric complex is a functional ATP-dependent helicase, we developed a ssDNA binding assay using fluorescently-labelled HaloTag-JF646-RPA (expressing RPA14, RPA70, and Halo tagged RPA32) calibrated against λ phage arms dsDNA containing 30-, 60- and 90-nt poly (dA) bubbles. As seen in Fig. 4F, RPA did not bind bubbleless dsDNA and RPA binding linearly increased for 30 to 90 nt ssDNA bubbles. RPA binding was then determined for mN-LT binding to Multi.Ori98 DNA under various conditions, supplemented with 1 mM dNTP. Trimeric mN-LT, which cannot form an active helicase, melts origin sequence through ATP-independent strand invasion^5^ and binds RPA equivalent to ∼30 nt ssDNA (Fig 4F, dotted line, S12). Both single-hexamer and double-hexamer LT (red bars) accumulations increase RPA binding to > 90 nt equivalents, consistent with ATP-dependent helicase unwinding. Since this assay does not account for LT foot-printing or potential primase dsDNA fill-in, RPA-binding intensity for LT-induced unwinding underestimates the actual ssDNA exposed in this assay, which could be greater than 90 nt. Notably, addition of sT (yellow bars) markedly increased the frequency of double-hexamer formation (Fig. 4A), which was reversed by addition of Ver-155008 (green bars), but the RPA-binding for 3-, 6- and 12-mer LT were similar. Substitution of ATP with non-hydrolyzable AMP-PNP (blue) or mutation to LT at Walker A (LT^K599R^, purple) or Walker B (LT^E640A/D641A^, magenta) sites that abolish ATP-dependent helicase activity generated ssDNA equivalent to trimeric wild-type LT. Helicase processivity, which requires assembly and movement of the fully-assembled and licensed cellular replisome on DNA^39^, was not seen this this assay.

### HSP70 Binds and Promotes Virus-Free Mouse MCM Complex Binding

To test whether HSP70 similarly promotes assembly of the cellular replicative helicase, we used biotinylated λ-DNA as a well-characterized naked dsDNA substrate, which is appropriate for these experiments because mammalian MCM loading is not governed by a short sequence-specific origin motif but instead occurs at thousands of distributed sites across the genome, making double-stranded DNA of any sequence a suitable substrate for single-molecule analysis of MCM–DNA interactions. The requirement for HSP70 in viral helicase assembly raised the possibility that this function extends to cellular replication machinery. To test this, mouse MCM complexes were assembled on biotinylated 48kbp λ phage DNA in the absence of viral proteins. Individual MCM subunits (mEmerald-MCM3, -MCM4 or -MCM6^40^) in U2OS SMADNE extracts co-localized with mScarlet-HSP70 in discrete foci on λ DNA (**Fig. 5A, S13**) . Direct binding of endogenous HSP70 to DNA-bound MCM complexes, was confirmed by antibody capture experiments. Emerald-MCM6 was first loaded onto λ DNA and then transferred to a C-Trap channel containing 1 μg/ml anti-HSP70-atto594 antibody (**Fig. 5B, S14**). Following transfer to a wash channel, persistent co-binding of both MCM6 and HSP70 antibody was observed. In contrast, when bound mEmerald-MCM6 was treated with 5 μM Ver-155008, mScarlet-HSP70 co-binding was lost (**Fig. 5C, S15**). To evaluate the impact of HSP70 inhibition on MCM assembly, six independent mEmerald-MCM6/λ DNA binding experiments were performed (**Fig. 5D**). MCM complex binding diminished (but was not eliminated) and binding times were reduced when treated with the HSP70 inhibitor, consistent with impaired formation or stabilization of functional MCM complexes in the absence of HSP70 activity.

**Fig 5.**
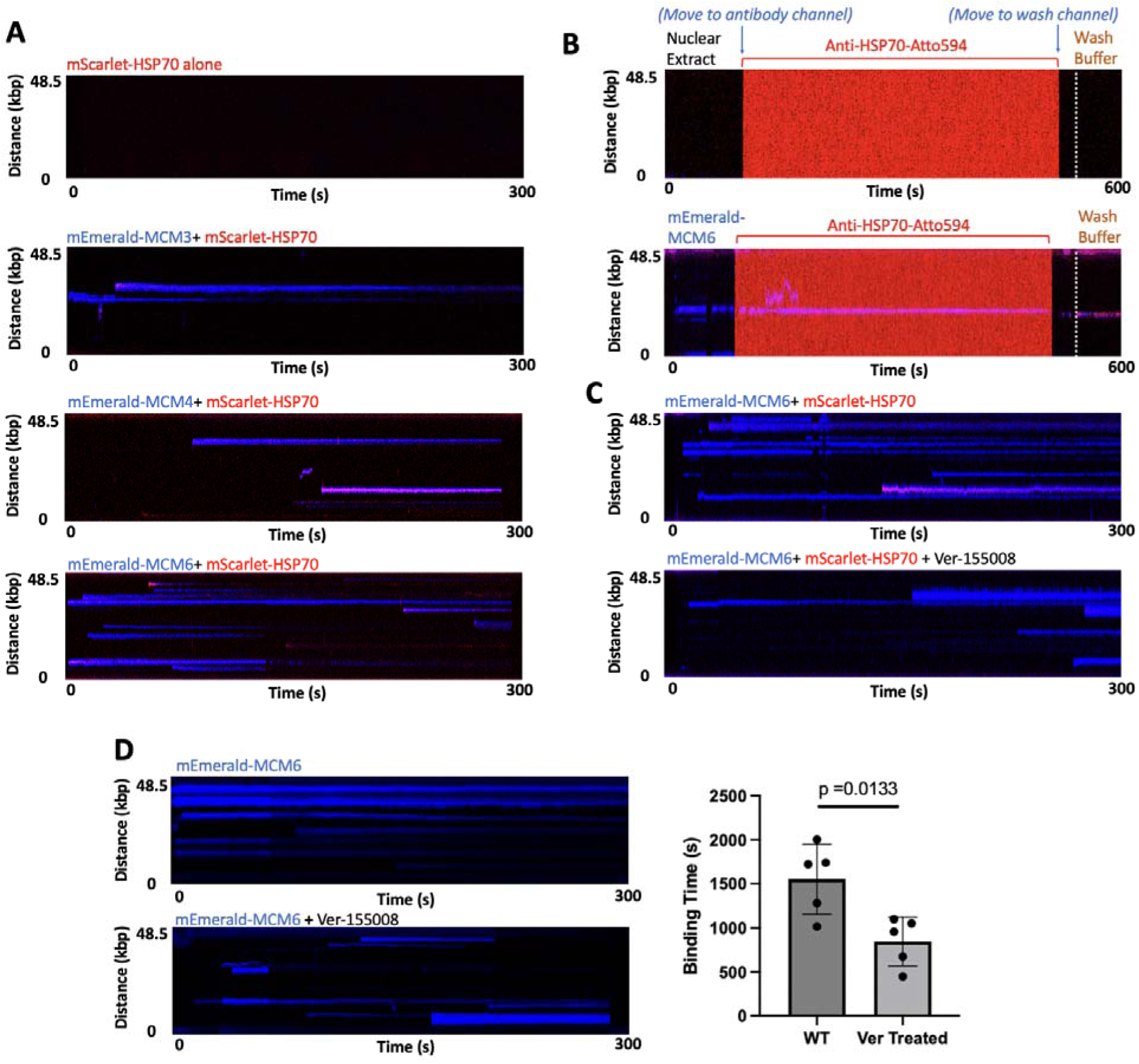
HSP70 inhibition reduces the efficiency of MCM binding to λ bacteriophage DNA. **(A)** MCM3, MCM4, and MCM6 interact with mScarlet–HSP70 on λ DNA. In C-Trap assays, nuclear extracts expressing fluorescent mEmerald–MCM3/4/6 (blue) were individually mixed with mScarlet–HSP70 (red) in the laminar-flow channel and applied to biotinylated λ DNA tethered between streptavidin-coated beads. mScarlet–HSP70 signal was detected at sites of MCM3, MCM4, and MCM6 binding. **(B)** MCM recruits endogenous HSP70 on λ DNA. Nuclear extract with or without mEmerald–MCM6 was flowed over λ DNA, then transferred to Anti-HSP70-Atto594 (1 µg/ml) and to antibody-free wash buffer. Antibody signal required bound MCM6 that was absent from MCM6-free extract. Both signals persisted and remained co-localized after washing (∼550–600 s). **(C)** HSP70 inhibition blocks recruitment of HSP70 to DNA-bound mEmerald–MCM6. Upper panel: mEmerald–MCM6 (blue) colocalizes with mScarlet–HSP70 (red) on λ DNA. Lower panel: Pretreatment of nuclear extracts with 5 µM Ver-155008 abolished mScarlet–HSP70 binding to mEmerald–MCM6. **(D)** HSP70 inhibition decreases mEmerald–MCM6 binding efficiency by ∼55%. Left: Pretreatment with 5 µM Ver-155008 reduced both the number of binding events and the dwell time of mEmerald–MCM6 on DNA. Right: Quantitation of total binding time per kymograph shows an approximate 55% reduction upon Ver-155008 treatment. Representative kymographs and data are recruited from 6 kymographs per condition.

## DISCUSSION

This study identifies a previously unrecognized, direct role for the MCV sT antigen in viral DNA replication. Our data establish that sT is an integral component of the viral replisome and functions directly in replication initiation by enabling HSP70-dependent remodeling of LT at the origin. Although LT alone can assemble into dodecameric helicases and melt origin DNA^5^, single-molecule C-Trap analysis reveals that sT significantly enhances higher-order LT assembly and replication competence through HSP70 recruitment and remodeling. The sT mediated increase in dodecameric LT frequency and complex lifetime on origin DNA, both abolished by HSP70 inhibition or DnaJ mutation, provides strong evidence for HSP70-dependent enhancement of replication-competent helicase assembly.

The assembled dodecamer is a functional helicase. Using RPA as a single-molecule reporter of exposed single-stranded DNA, we find that unwinding at the origin separates into two components: a hydrolysis-independent melting of approximately 45 nt, which persists when ATP hydrolysis is blocked by AMP-PNP or by Walker A or Walker B mutation, and a substantially larger hydrolysis-dependent extension that increases with assembly state and is maximal at the dodecamer. The first corresponds to the strand-invasion melting we described previously, which proceeds without ATP hydrolysis and without double-hexamer formation. The second is helicase activity, and it is the activity whose assembly sT and HSP70 promote. Separating the two resolves a persistent ambiguity in interpreting origin-bound LT: the appearance of single-stranded DNA is not by itself a signature of helicase function.

In contrast to LT, which engages origin DNA through its sequence-specific origin binding domain and maintains a fixed position at the Ori98 sequence^6^, sT alone displays only non-specific DNA association, migrating freely under applied shear force and lacking a defined origin contact site (**Fig. S2, S3**). Within the LT-HSP70-sT complex, however, sT adopts a stable, origin-proximal position adjacent to but not overlapping with LT, tethered in place through its HSP70-mediated connection to origin-bound LT. The observed spatial separation between LT and sT fluorescence signals on DNA therefore reflects this architectural arrangement with LT anchored at its sequence-specific binding site and sT positioned within its diffusion-limited range through the HSP70 bridge, rather than a precise rigid molecular span. This is consistent with the dynamic positional behavior documented for protein complexes studied by single-molecule imaging on DNA^34,35^. This arrangement also depends on intact DnaJ domains in both LT and sT and reflects the formation of an ordered LT-HSP70-HSP70-sT assembly rather than a passive co-localization event. The collapse of this spatial organization upon HSP70 inhibition or DnaJ domain mutation underscores the architectural contribution of HSP70 to replisome formation.

These data also reveal an unexpected role for HSP70 as a structural and remodeling factor within a eukaryotic viral replication complex. Unlike its canonical chaperone activity, in which HSP70-HSP40 recycles after substrate remodeling^41^, HSP70 remains stably associated with the LT-sT complex and constitutes a core component of the MCV replisome (**Fig. 6**). Previous work by Campbell et al.^28^ demonstrated that the DnaJ domain of SV40 large T antigen is required for efficient DNA replication of the related polyomavirus SV40, and it was proposed that HSP70-mediated rearrangement of LT complexes underlies this requirement — a mechanistic framework that our single-molecule data now directly confirm and extend to MCV, where sT serves as an obligate co-chaperone adaptor. This mechanism parallels, but is distinct from, the activity of the DnaK-DnaJ system during replisome remodelling at the origin of bacteriophage DNA replication in E. coli ^2,42^. The DnaK-DnaJ ensemble, together with the GrpE nucleotide exchange factor, activates the DnaB helicase to initiate DNA replication. Notably, GrpE is dispensable for DNA replication in vitro when DnaK is present at high levels ^43^.

**Fig 6.**
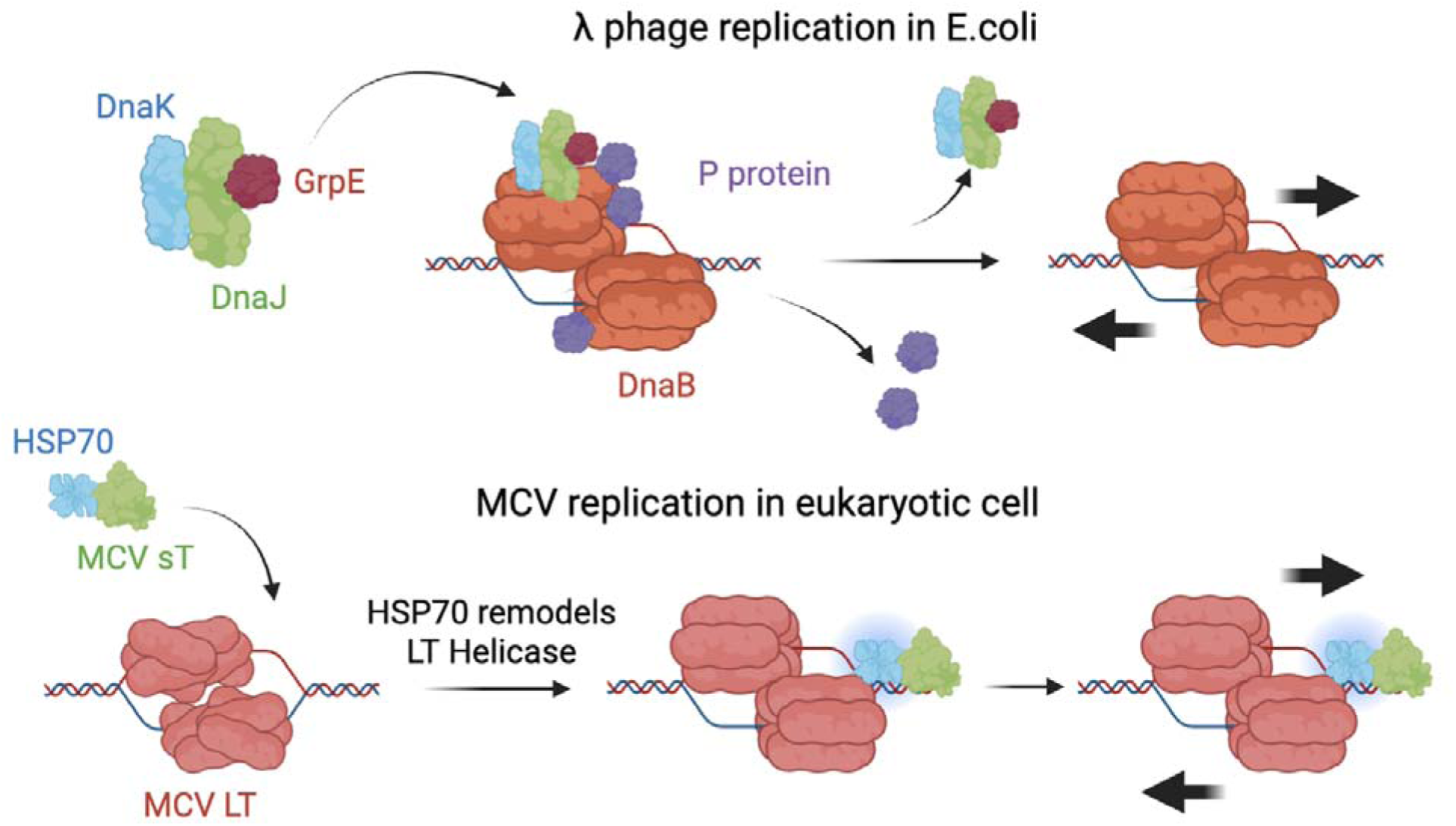
MCV small T antigen (sT) functionally mimics bacterial DnaJ to recruit HSP70 and facilitates remodeling assembly of the viral replication complex. In E. coli, DnaJ recruits the HSP70 homolog DnaK and nucleotide exchange factor GrpE to displace the inhibitory P protein from the helicase DnaB, thereby activating λ phage DNA replication. In eukaryotic cells, Merkel cell polyomavirus (MCV) sT employs its DnaJ domain to recruit host HSP70 and promote remodeling of large T antigen (LT) into a functional helicase on the viral origin of replication.

Intriguingly, Lin et al.^44^ report that HSP70-HSP40 participates in human papillomavirus (HPV) replication, despite the absence of DnaJ motifs in the HPV E1 and E2 replication proteins. This raises the possibility that HSP70-mediated remodelling could be a conserved eukaryotic, as well as prokaryotic, replication mechanism. Extending this paradigm, we demonstrate that mammalian MCM, the cellular replicative helicase corresponding to LT, also associates with HSP70 when bound to DNA. HSP70 inhibition reduced, but did not abolish, MCM binding suggesting that HSP70 is not strictly required for initiation but substantially enhances efficient double-hexamer formation. Given the structural and mechanistic parallels between polyomavirus LT and the cellular MCM replicative helicase^45^, our findings further motivate re-evaluation of how HSP70 and related chaperone networks regulate host replication initiation and replisome maturation.

Pharmacologic dissection of HSP70 function reinforces its central role in MCV replication. Inhibitors targeting ATP binding, DnaJ-mediated activation, or nucleotide exchange each disrupted the LT-sT architecture and prevented the formation and maintenance of replication-competent LT dodecamers. Although HSP70 inhibitors have been examined in MCV-positive Merkel cell carcinoma, in which the LT protein is mutated and replication-inactive^46^, their effects were previously attributed to general roles in protein folding, stabilization, and trafficking ^47,48^. Our data instead place HSP70 at the core of the MCV replisome architecture, and demonstrate that, in the context of replicating MCV, HSP70 activity is mechanistically linked to the assembly of the origin-bound helicase itself.

While our data define a direct role of sT- and HSP70-dependent remodeling of LT, several limitations remain. Our single-molecule experiments used nuclear extracts rather than fully purified systems, leaving open whether additional host cofactors contribute to LT-HSP70-sT assembly. Our measurements likewise establish ATP-hydrolysis-dependent unwinding at the origin but do not address processivity: sustained replication-fork unwinding requires assembly and licensing of the wider human replisome, which we have not reconstituted on the C-Trap, and we make no claim of processive unwinding here. Moreover, the structural transitions within LT induced by HSP70 remain undefined and will require high-resolution approaches such as cryo-EM. Although HSP70 enhances mammalian MCM binding and stability on DNA, these assays do not fully capture the complexity of cell-cycle-regulated replisome assembly in vivo, and the order of LT-HSP70-sT assembly, which may occur prior to DNA binding, will need to be established. Addressing these points will clarify the generality of chaperone-mediated helicase remodeling in eukaryotic DNA replication.

The central observation nonetheless holds across approaches: a viral protein recruits a host chaperone to convert an inefficiently assembling helicase into a replication-competent one, and the resulting complex unwinds origin DNA in a manner that requires ATP hydrolysis. That a human tumor virus should depend on the same chaperone-mediated strategy as bacteriophage λ suggests that remodeling of replication initiation complexes by HSP70-family chaperones is more broadly distributed than the division between prokaryotic and eukaryotic replication would imply.

## METHODS

### Cell lines

293 cells (ATCC) were maintained in Dulbecco’s Modified Eagle Medium (ThermoFisher) supplemented with 10% fetal bovine serum (FBS). U2OS cells were cultured in McCoy’s 5A medium (ThermoFisher) with 10% FBS. UISO cells were grown in RPMI 1640 medium (ThermoFisher) supplemented with 10% FBS, sodium pyruvate (ThermoFisher), and non-essential amino acids (ThermoFisher). All cell lines were maintained at 37°C in a humidified 5% CO₂ atmosphere.

### Plasmid Construction and Mutagenesis

Codon-optimized MCV LT and sT sequences were cloned into parental vectors bearing mNeonGreen, mScarlet, or HaloTag via standard restriction-ligation. Point mutations (LT^D44N^ and sT^D44N^) were introduced using the QuickChange Lightning Site-Directed Mutagenesis Kit (Agilent) following the manufacturer’s protocol.

### Origin Replication Assay

293 or U2OS cells were seeded in 6-well plates and transfected with 1 µg total plasmid DNA using Lipofectamine 2000 (ThermoFisher). After 48 h, genomic DNA was isolated using the DNeasy Blood and Tissue Kit (Qiagen). To eliminate unreplicated bacterial DNA and linearize replicated Ori98 DNA, 1.25 µg of DNA was digested overnight with BamHI and DpnI.

### Quantification of Replication by qPCR

Digested DNA (5 ng) was quantified by qPCR using PowerUp™ SYBR™ Green Master Mix (ThermoFisher) on a CFX Duet Real-Time PCR Machine (Bio-Rad).

- Ori98 primers: Fw: 5’-GCCGCCAAGGATCTGATG-3’ Rev: 5’-CTGCGCAAGGAACGCCCGTCG-3’.
- GAPDH normalization primers Fw: 5’-TGTGTCCCTCAATATGGTCCTGT-C-3’ Rev: 5’-ATGGTGGTGAAGACGCCAGT-3’).

Relative replication levels were calculated using the ΔΔCt method.

### Immunoblotting

Transfected cells were lysed in RIPA buffer (150 mM NaCl, 1% NP-40, 0.5% DOC, 0.1% SDS, 50 mM Tris-HCl pH 7.4) supplemented with protease inhibitors (0.2 mM sodium orthovanadate, 0.3 mM PMSF, and 1 ug/mL each of leupeptin, pepstatin A, and aprotinin). Lysates were sonicated on ice (20% amplitude, 5 s pulses x4) using a Fisherbrand™ Model 505 Sonic Dismembrator (ThermoFisher). Samples were mixed with 5x Laemmli buffer, resolved by SDS-PAGE, and transferred to nitrocellulose. Membranes were probed overnight at 4°C with primary mouse monoclonal antibody against MCV LT (CM2B4, 1:500), MCV sT (2T2, 1:200), or HSP70 (5A5, 1:1000). After washing, blots were incubated at room temperature with IRD800-conjugated goat anti-mouse secondary antibody (LI-COR Biotechnology, 1:10000) and Rhodamine-conjugated anti-tubulin antibody (Bio-Rad, 1:10000). Protein bands were visualized using the ChemiDoc™ MP Imaging System (Bio-Rad).

### Co-immunoprecipitation

U2OS cells in 6 well plates were co-transfected with 0.75 µg each of LT, sT, empty vector, and Flag-HSP70 plasmids using Lipofectamine 2000 (ThermoFisher). After 48 h, cells were lysed with 500 µl of lysis buffer (2% Triton, 5mM NaF, 5% glycerol, in PBS buffer, supplemented with PMSF(Sigma)). Lysates were incubated overnight at 4°C with the anti-Flag antibody (Sigma, F3165, 1:1000), followed by incubation with Dynabeads™ Protein G (Invitrogen, 40 µl) for 2 h. Beads were washed three times with lysis buffer, boiled in SDS loading buffer for 10 min, and analyzed by SDS-PAGE and immunoblotting as described above.

### Nuclear Extract for Single-Molecule Assays

To prepare nuclear extract for a protein, UISO cells in 6-well plates were transfected with 2 µg of the corresponding plasmid and 2 µl of Lipofectamine 2000. After 48 h, nuclear extracts were prepared using the NE-PER™ kit (ThermoFisher) according to the provided protocol and diluted 1:100 in C-Trap reaction buffer (25 mN Tris–acetate, 10 mM magnesium acetate, 50 mM potassium acetate, 0.1 mM EDTA, 1 mM ATP, and 3 mM DTT) at 1:100 ratio immediately before use. This includes untagged LT, untagged sT, mN-LT, LT-mS, EGFP-sT, mN-sT, EGFP-HSP70, mS-HSP70, mEmerald-MCM3, mEmerald-MCM4, and mEmerald-MCM6. For HaloTag proteins, an additional labeling step was performed by adding one vial of Janelia Fluor® HaloTag® ligand (549, 635, or 646; Promega) into each well of a 6-well plate 30 min before harvest. This includes HaloTag-JF549-LT, HaloTag-JF646-LT, HaloTag-JF549-sT, and HaloTag-JF646-sT.

For HSP70-inhibitor studies, diluted nuclear extracts were incubated with 5 µM VER-155008, 50 µM MAL3-101, or 50 µM MKT-077 for 10 min at RT prior to C-Trap analysis.

### Biotinylated DNA Substrates

pMC.Ori98 was linearized with XmaI and EcoRI-HF, column purified with NucleoSpin Gel and PCR Clean-up kit (Macherey-Nagel, 740609.50), self-ligated with T4 DNA ligase, then end-labelled with biotin-14-dCTP (AAT Bioquest, 17019) and biotin-11-dGTP (AAT Bioquest, 17015) using Klenow Fragment (NEB, M0210S). After column purification, biotinylated DNA was diluted 1:200 in PBS prior to use. For MCM binding experiments, biotinylated λ-DNA (48.5 kbp, LUMICKS) was used as the DNA substrate, diluted 1:200 in PBS prior to use. λ-DNA was chosen as a well-characterized long dsDNA molecule suitable for single-molecule analysis of non-sequence-specific DNA-binding proteins, consistent with published single-molecule studies of eukaryotic replication-associated proteins^29^.

### Optical Tweezer-Fluorescence Microscope (C-Trap)

Single-molecule assays were performed on a LUMICKS C-Trap instrument with dual optical traps. Streptavidin-coated polystyrene beads (1:100 dilution from 0.5% w/v concentration) were captured with dual optical tweezers in channel 1, then moved to channel 2 to capture a single biotinylated dsDNA molecule. The captured DNA were then moved to channel 4 containing nuclear extracts with fluorescently labeled proteins. Binding events of mNeonGreen-, mScarlet-, or HaloTag bound Janelia Fluor® 549 or 646 ligands were visualized using the integrated confocal fluorescence microscope. Kymographs were recorded along the DNA in 2D, with at least 300s duration.

### Image display and intensity quantification

All kymographs within a given comparison were acquired with identical laser powers, pixel dwell times and detector settings. For display, panels compared with one another — the two conditions in Fig. 1F, the individual fluorescence channels of a single trace in Figs. 3D–G, 5B and S7, and all panels within a figure — were rendered with the same look-up table (RGB for merged images and pseudocolor red, green, or blue for split images) and the same intensity range (8-bit, display range: 0-255). No panel-specific contrast adjustment was applied at any stage. Quantitative measurements on event intensity and multimer level determination were made on intensity observed from the ImageJ files. FRET signal quantitation (Fig. S11) was made on raw photon counts extracted from the .h5 acquisition files using LUMICKS Lakeview software, not on rendered images.

### Data Analysis

Fluorescence trajectories were extracted using LUMICKS Lakeview software. The extracted data (in .h5 format) were analyzed for binding durations, positions, and photon counts. Kymographs were rendered in ImageJ, applying the look-up tables and display ranges specified above. Cross-sectional intensity profiles used to determine binding-event positions were quantified using the “plot profile” function on ImageJ. Gaussian fitting was applied to determine each binding event location on the DNA, where the center of each fluorophore signal peak was located. The distance between peaks were calculated, using length of the DNA as a reference.

For statistical analysis of LT, sT, and HSP70 binding dynamics, including *k*on, binding lifetime, binding efficiency, and multimerization level, only stable binding events lasting >5 s were included. Events were analyzed from at least 6 independent kymographs. For LT multimerization analysis, we applied the Hidden Markov Model analysis to kymographs, as previously described^5^.

For localization analysis, LT–sT or LT–HSP70–sT co-bound complexes were selected from kymographs based on stable co-binding events lasting >5 s. Cross-sectional fluorescence intensity profiles were generated at least 5 s after the initial binding event to ensure that the complex had reached a stable binding state.

### Association Constant kon Determination

Binding of mNeonGreen-LT dodecamers to Ori98 was modelled as

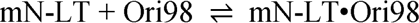

Concentration of the mN-LT•Ori98 complex:

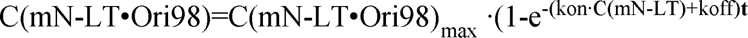

**Fig. 2A** depicts:

mN-LT dodecamer binding to Ori98 persists over 1000s, thus Koff ≈ 0

Therefore

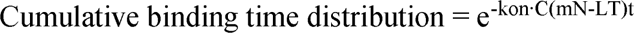

C(mN-LT)is maintained as constant because mN-LT nuclear extract was refreshed continuously in the flow cell.

The exponential constant of the curves K = −kon C(mN − LT) is proportional to kon. Comparing K of mN-LT versus mN-LT+sT reveals a 50% increment in kon for mN-LT binding to Ori98 with the presence of sT.

### Single-molecule FRET measured by acceptor photobleaching

Single-molecule Förster resonance energy transfer (FRET) measurements were performed using the C-Trap with mN-LT as the donor and HaloTag–JF549-sT as the acceptor. Three experimental conditions were examined: wild-type LT with wild-type sT, LT^D44N^ with wild-type sT, and wild-type LT with sT^D44N^.

FRET was assessed by acceptor photobleaching followed by donor fluorescence dequenching. Initially, the donor was excited using a 488 nm laser at 5% power, and fluorescence signals from both mN and JF549 were recorded to establish FRET intensity. The 488 nm laser was then switched off, and a 561 nm laser at 50% power was applied to completely photobleach the JF549 acceptor. Following acceptor photobleaching, the 561 nm laser was switched off, and 488 nm excitation at 5% power was resumed to monitor the recovery of mN donor fluorescence resulting from the loss of FRET-mediated energy transfer. Fluorescence intensities before and after acceptor photobleaching were quantified using the photon count. An increase in donor fluorescence following acceptor photobleaching was interpreted as evidence of FRET between mN and JF549, indicating close spatial proximity between the two fluorophores.

The photon count per second were recorded during all three phases from 6 kymographs for each condition. The FRET efficiency E was calculated as:

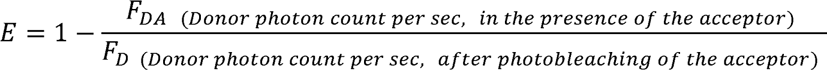

### Quantitation of MCV large T antigen-mediated DNA melting using HaloTag–JF646–RPA Preparation of poly(dA) bubble DNA standards

Synthetic DNA substrates containing defined single-stranded poly(dA) bubbles of 30, 60, or 90 nucleotides were generated by annealing complementary oligonucleotides containing central poly(dA) stretches flanked by complementary double-stranded DNA regions (Table .S2). Following annealing, the resulting DNA substrates contained a central single-stranded bubble of defined length, mimicking the locally unwound DNA intermediate generated during replication initiation. The bubble-containing substrates were ligated between two 6.3-kb biotinylated DNA handles using the DNA tethering kit (LUMICKS), generating DNA constructs suitable for C-Trap measurement.

### Calibration of Halo tagged RPA32 fluorescence intensity using poly(dA) bubbles

DNA constructs containing 30-, 60-, or 90-nucleotide poly(dA) bubbles were introduced into the C-Trap system and tethered between two streptavidin-coated beads through the biotinylated DNA handles. HaloTag–JF646–RPA containing nuclear extracts were introduced into the flow channel to allow RPA binding to the exposed single-stranded DNA regions. Fluorescence images were acquired under consistent imaging conditions, and the maximum fluorescence intensity of RPA associated with each DNA bubble was quantified following background subtraction. The measured fluorescence intensities were plotted against the corresponding known bubble lengths (30, 60, and 90 nt), and linear regression was performed to generate a calibration curve relating HaloTag–JF646–RPA fluorescence intensity to the amount of exposed single-stranded DNA. The resulting calibration equation was subsequently used to estimate the extent of DNA melting in LT-containing experiments.

### Measurement of LT-mediated DNA melting

DNA melting by mN-LT was examined using the C-Trap system under the indicated experimental conditions. LT-containing nuclear extracts were introduced together with HaloTag–JF646–RPA containing nuclear extracts, with the same amount of RPA-containing extract maintained across all experimental conditions. Experiments were performed in the C-Trap reaction buffer described above, supplemented with 0.1 mM dNTPs to support nucleotide-dependent DNA synthesis. LT-mediated DNA unwinding was monitored through the recruitment and accumulation of HaloTag–JF646–RPA on newly exposed single-stranded DNA regions. Fluorescence images were acquired using the same imaging settings as those used for the poly(dA) bubble calibration. The maximum intensity of HaloTag–JF646–RPA was measured and compared to the calibration intensity for an estimation of unwound ssDNA length.

## Data Availability

Source data underlying all quantitative display items in the main figures and supplementary figures, including binding efficiency measurements, lifetime distributions, multimerization frequencies, replication assay values, distance measurements, and FRET quantification, are provided as a Source Data file accompanying this paper. Kymograph imaging data supporting the findings of this study are available from the corresponding authors upon reasonable request. Source Data are provided with this paper.

## Code Availability

Fluorescence trajectory extraction was performed using LUMICKS Lakeview software (commercial). Kymograph processing and intensity quantification were performed using ImageJ (freely available at https://imagej.nih.gov). Hidden Markov Model analysis of LT multimerization was performed as previously described⁵. No custom code was generated for this study.

## Acknowledgments

We thank all members of the Chang-Moore Lab for their insightful discussions. We are particularly grateful to Bennett Van Houten and Matt Shaich for their valuable guidance on C-Trap analysis and performance, to Mashiro Shuda for experimental advice, and to Keith Jarrett and Carla Bley for their assistance with the manuscript.

## Author contributions

Conceptualization: PSM, YC

Methodology: PSM, YC, LW, JLB, PW

Investigation: LW, PV, XL

Funding acquisition: PSM, YC

Supervision: PSM, YC

Writing – original draft: PSM, YC, LW

Writing – review & editing: PSM, YC, LW, PV, XL, JLB, PW

## Supplementary Materials

Figs. S1 to S15

Tables. S2 to S2

