## Supplement for "A Human Tumor Virus Recruits HSP70 Chaperone to Remodel Its Replisome on DNA"

Li Wan^1^, Xiaochen Li^2,1^, Paul Vittot^1^, Jeffrey L. Brodsky^3^, Peter Wipf^4^, Patrick S. Moore^1,^*, Yuan Chang^1,^*

^1^Cancer Virology Program, Hillman Cancer Center, University of Pittsburgh, PA 15232, USA

^2^School of Medicine, Tsinghua University, Beijing, 100084, China

^3^Department of Biological Sciences, University of Pittsburgh, PA 15260, USA

^4^Departments of Chemistry, Pharmaceutical Sciences and Bioengineering, University of Pittsburgh, PA 15260, USA

**The PDF file includes:**

Figs. S1 to S15

Tables. S1 to S2

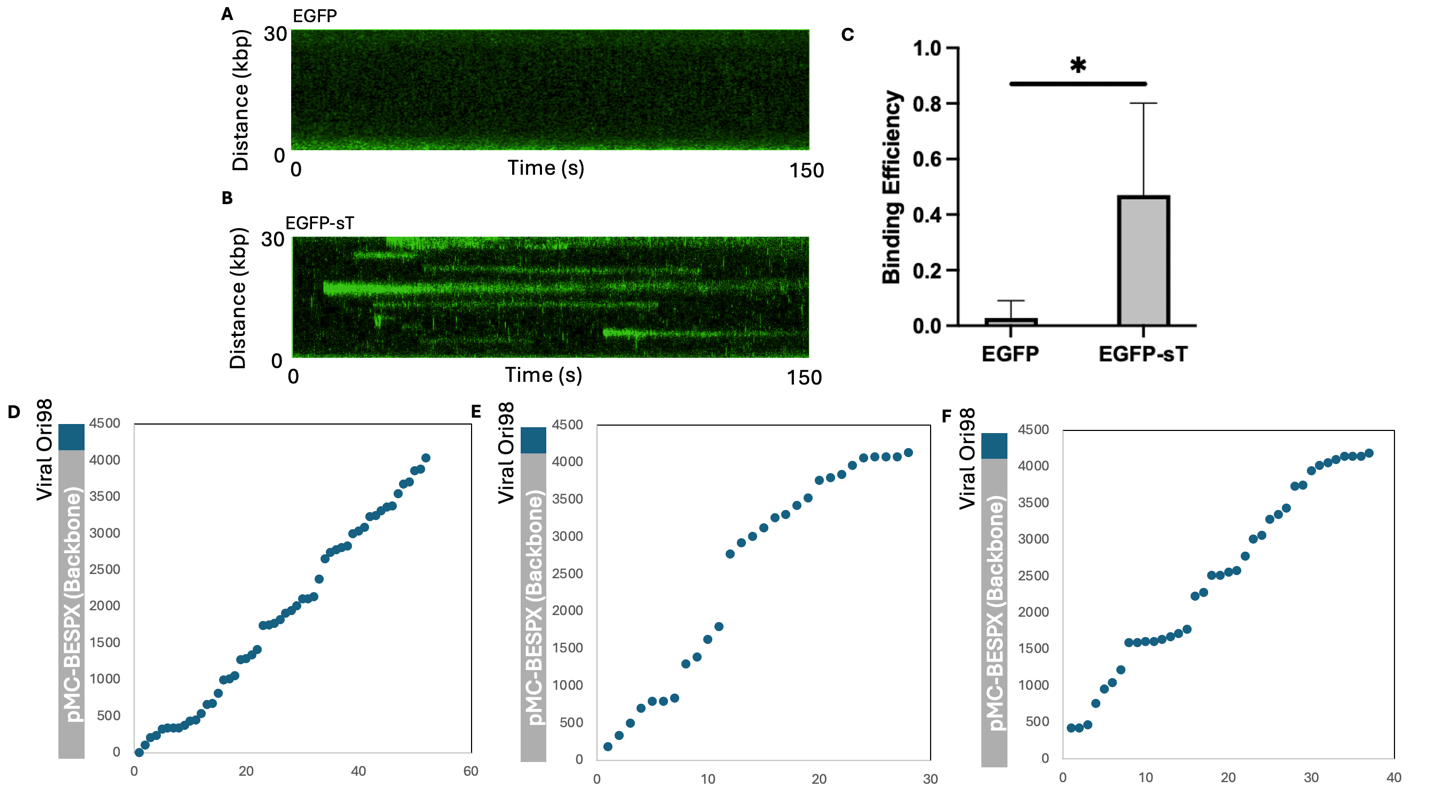

**Fig. S1.** sT binds non-specifically to DNA. (**A**) EGFP alone does not bind Multi.Ori98 DNA, whereas (**B**) EGFP-sT binds Multi.Ori98 DNA in a non-specific manner. (**C**) Quantitation of EGFP-sT binding efficiency to Multi.Ori98 DNA. (**D**) Distribution of EGFP-sT binding sites across Multi.Ori98 indicates a non-specific binding pattern. (**E**) Distribution of LT-mS binding sites when co-expressed with EGFP-sT, with enhanced binding to the viral Ori98. (**F**) Distribution of EGFP-sT binding sites in the presence of LT-mS, demonstrating enrichment to Ori98 sites due to recruitment of LT-mS.

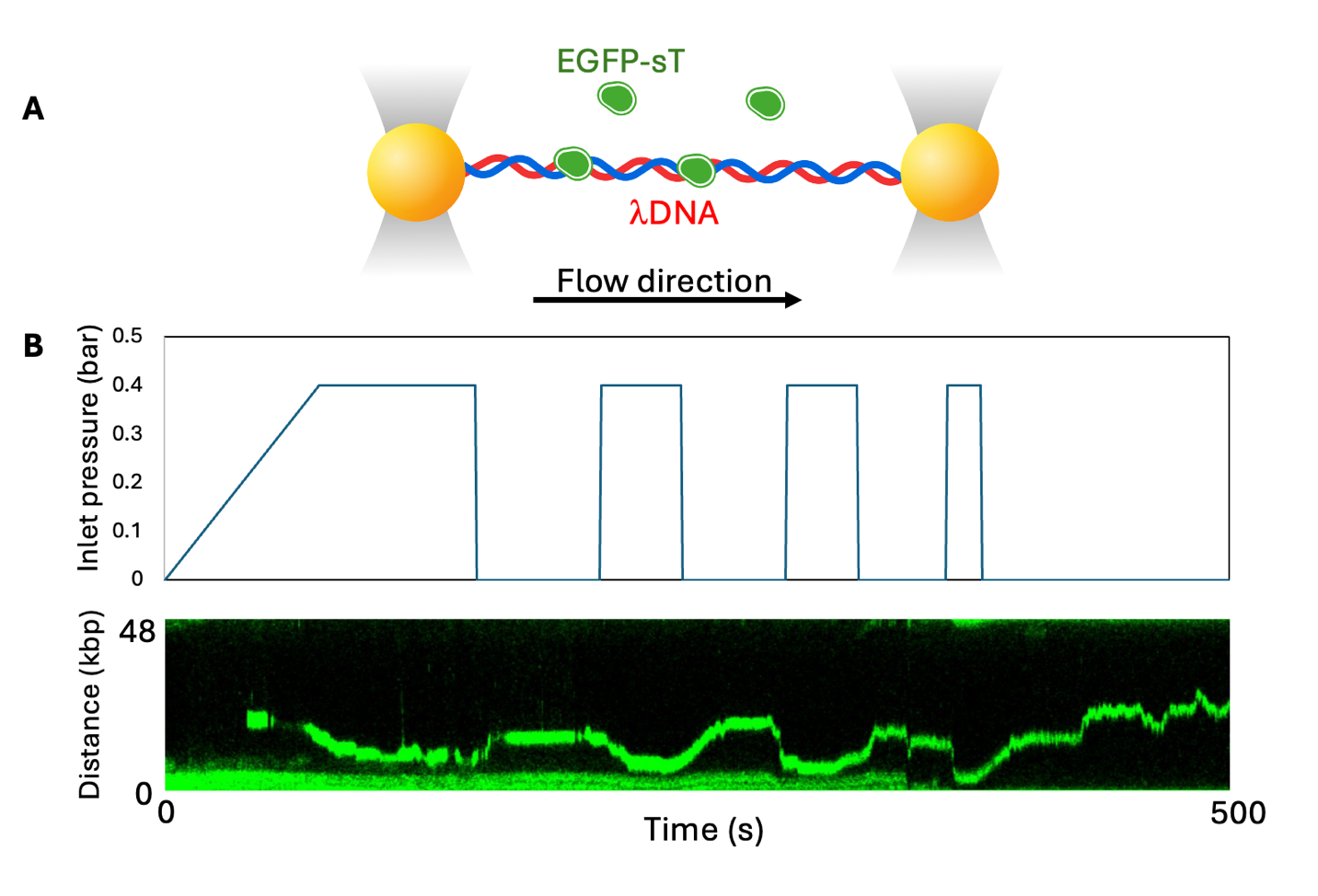

**Fig. S2. sT migrates along DNA under shear force.** (**A**) Experimental setup: Laminar flow was applied parallel to lDNA, generating shear stress on EGFP-sT bound to DNA. (**B**) EGFP-sT bound and migrated on λDNA (48.5 kb) under shear stress. Top: Inlet pressure was adjusted manually to create parallel shear stress to EGFP-sT on DNA. Bottom: Kymopgraph shows sT movement on DNA due to changes in shear stress. Increasing inlet flow pressure to 0.4 bar induced EGFP-sT migration, while releasing the pressure caused EGFP-sT movement in the opposite direction. This cycle was repeated multiple times.

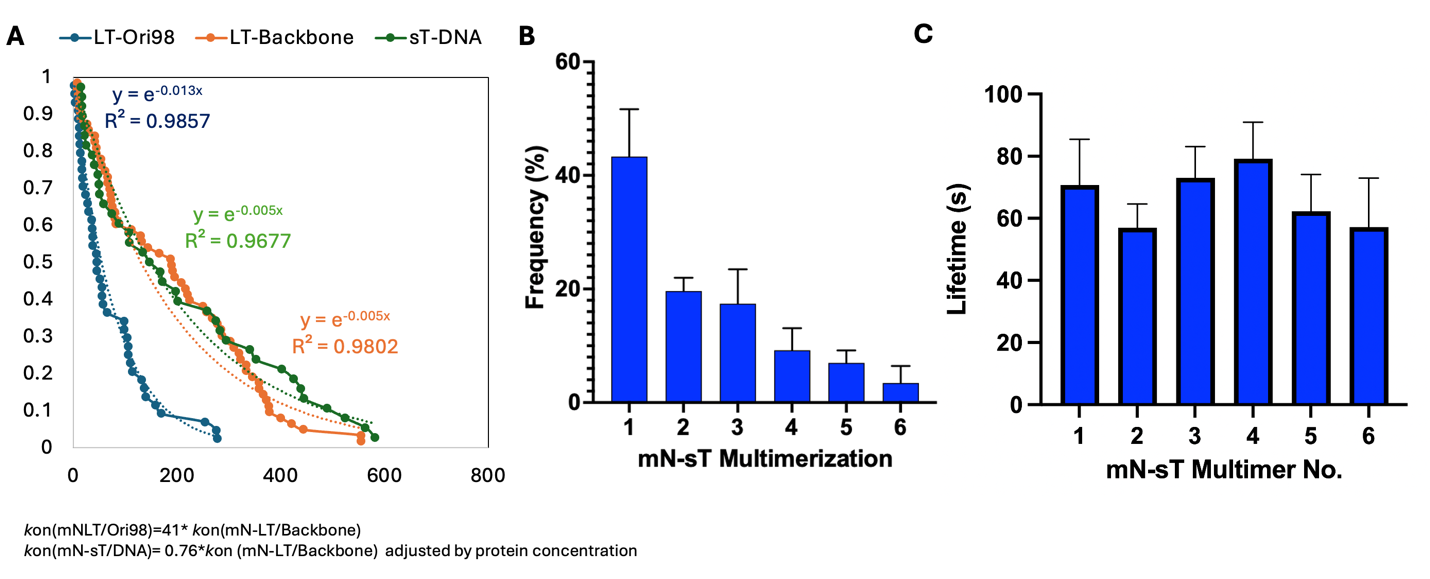

**Fig. S3. MCV sT–DNA binding dynamics.** (**A**) The *k*on for nonspecific sT binding to DNA is lower than that for nonspecific LT binding to the DNA backbone and lower than LT binding to Ori98. (**B**) mN-sT forms multimers of up to hexamer size on DNA, but no stabilized sT multimeric state was observed. (**C**) The DNA-binding lifetime of mN-sT is similar across different sT multimeric states and is shorter than that of the LT dodecamer (~200 s).

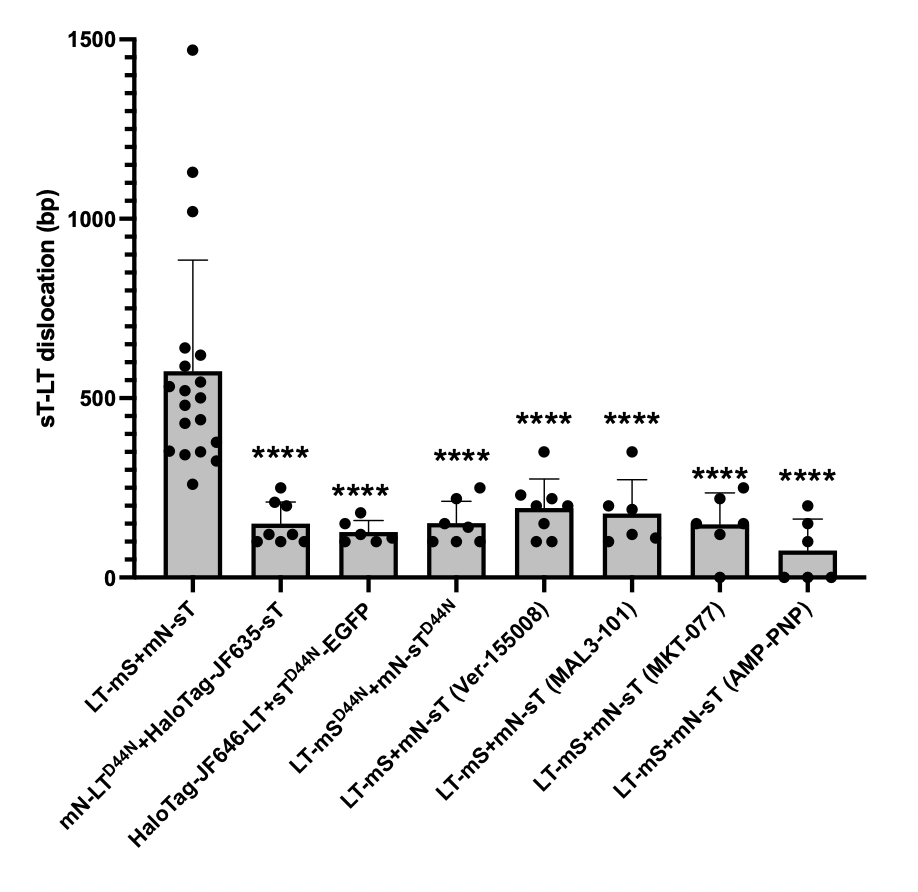

**Fig. S4**. **Distance between LT/LT^D44N^ and sT/sT^D44N^ binding locations on DNA, with and without HSP70 inhibitors.** Mutation of the DnaJ domain in either LT or sT led to collapse of the LT–sT spatial separation, indicating that HSP70 interactions with both LT and sT are essential for maintaining the gap. Inhibition of HSP70 using VER-155008 (5 µM), MAL3-101 (50 µM), or MKT-077 (50 µM) similarly resulted in collapse of the LT–sT spacing, further supporting the requirement for functional HSP70. Additionally, depletion of ATP using apyrase followed by supplementation with the non-hydrolyzable analog 1 mM AMP-PNP also caused LT–sT collapse, suggesting that ATP hydrolysis by HSP70 is necessary to maintain LT–sT spatial separation.

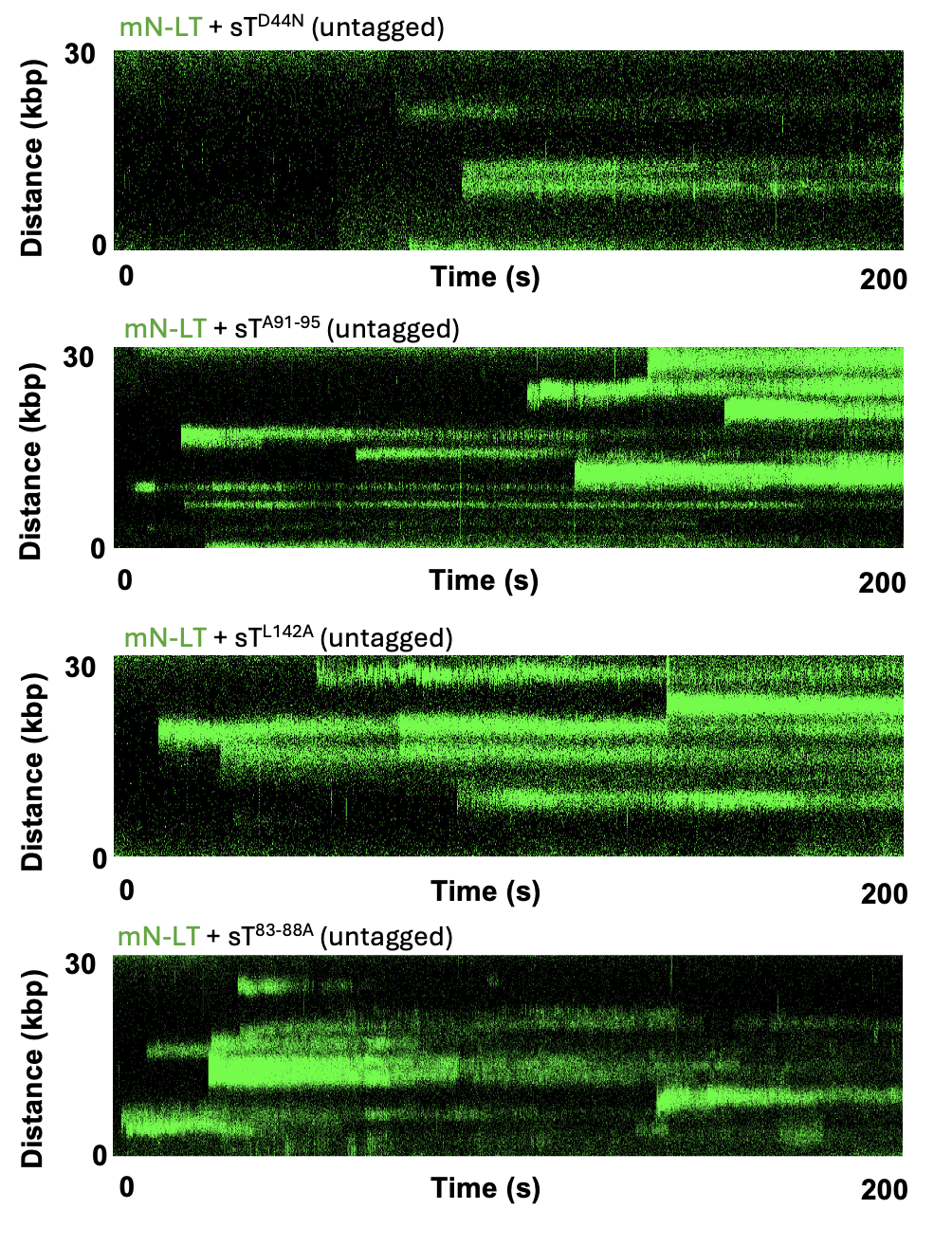

**Fig. S5. Representative kymographs of mN-LT binding to DNA in the presence of sT mutants**, including DnaJ mutant (D44N), LTSD (91–95A), PP2A-binding mutant (L142A), and EP400 mutant (sT83–88A). These kymographs correspond to the quantitative analysis of LT-origin binding efficiency presented in Fig. 2C. Representative from 6 kymographs per condition.

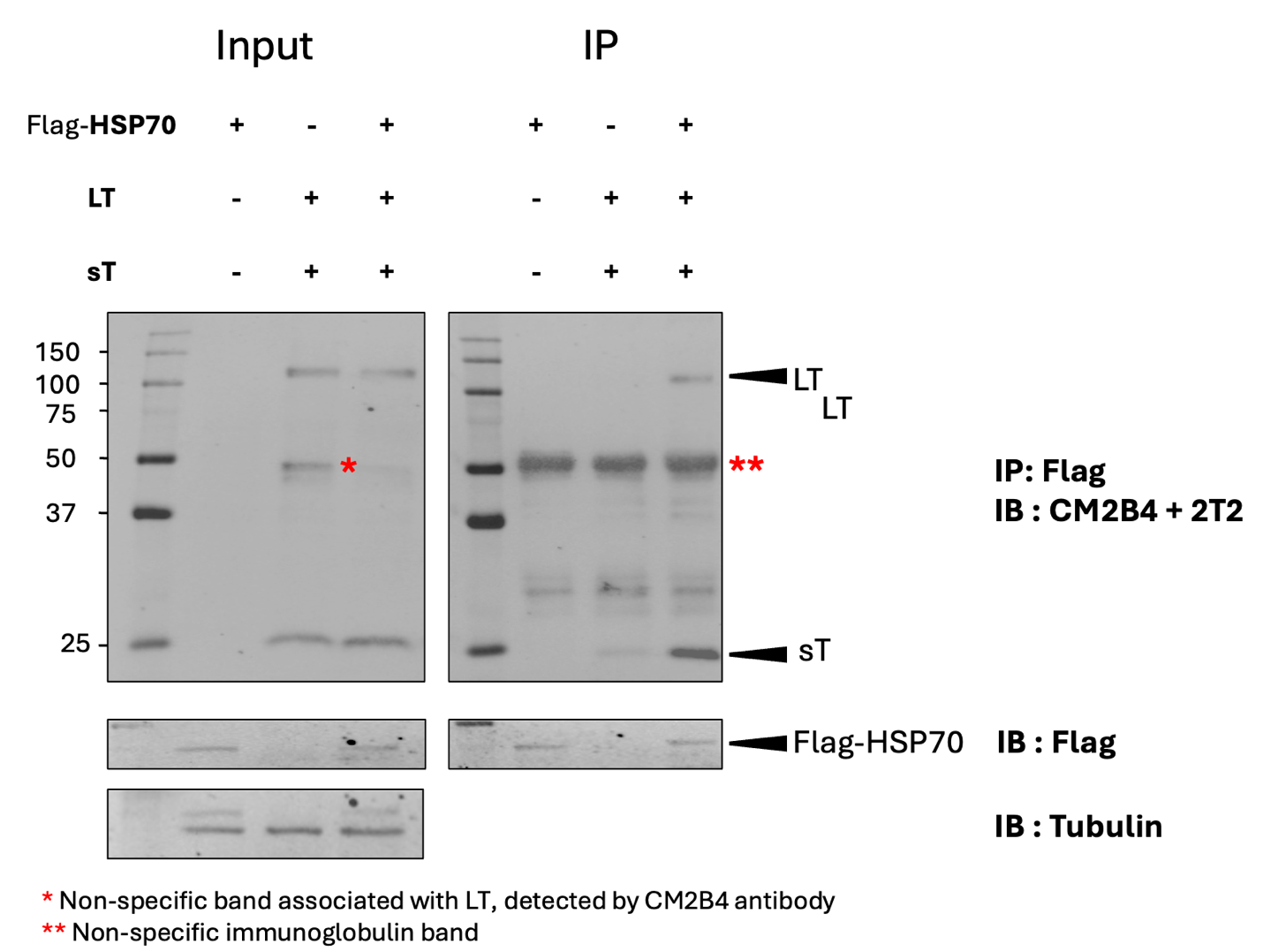

**Fig. S6. Co-immunoprecipitation of LT and sT with Flag-HSP70.** U2OS cells were co-transfected with LT and sT, or empty vector together with Flag-HSP70 plasmids for 48 hours, followed by immunoprecipitation using a Flag antibody. Co-expression with Flag-HSP70 enhanced the recovery of both LT and sT, indicating their association with HSP70.

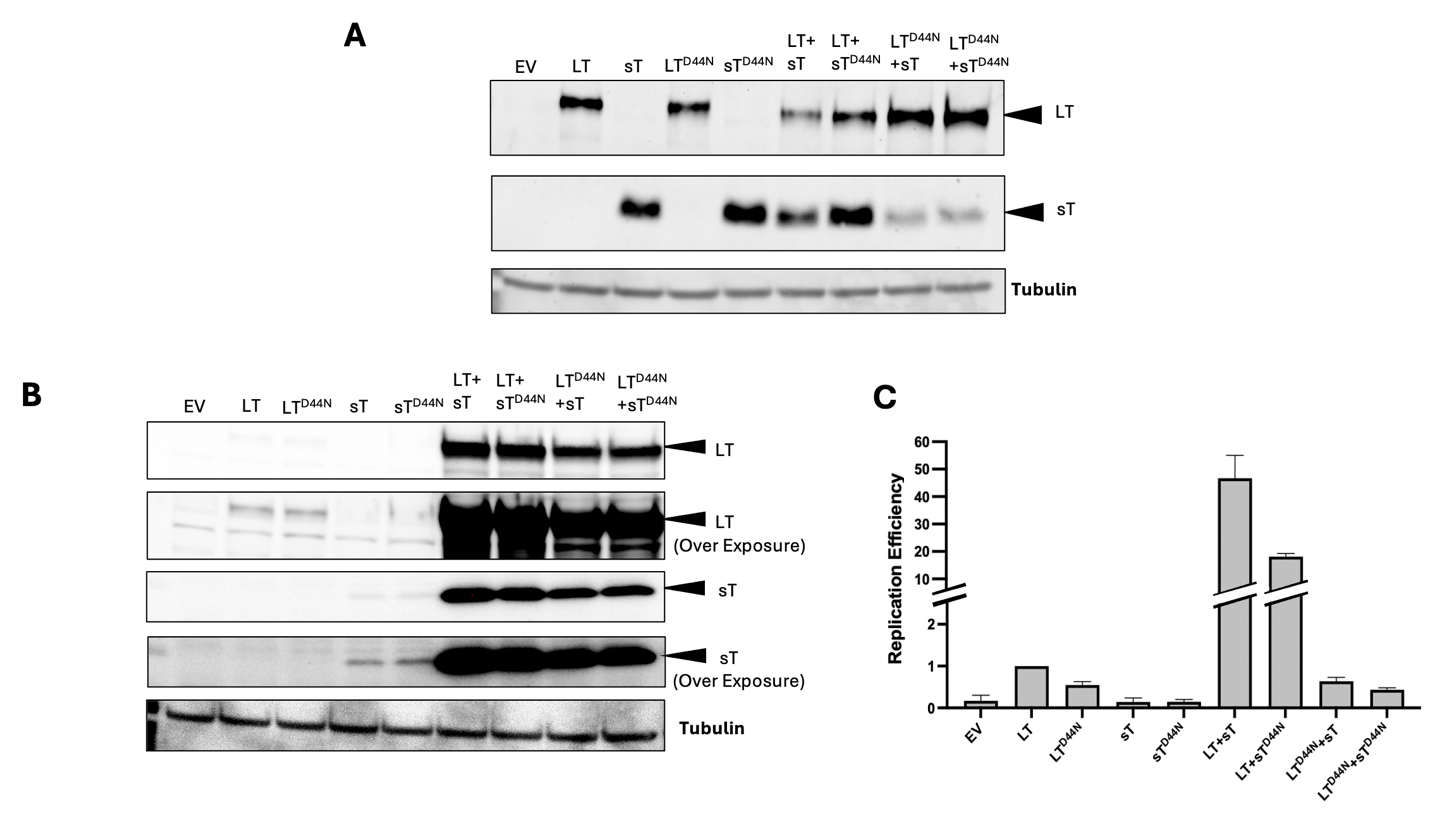

**Fig. S7. Mutations in the DnaJ domain of LT and/or sT reduce viral replication efficiency in U2OS and 293 cells.** U2OS and 293 cells were co-transfected with a viral origin containing plasmid and LT/LT^D44N^ + sT/sT^D44N^ for 48 h, then harvested and DNA purified to determine antigen expression level and viral origin containing plasmid replication efficiency. (**A**) Western blot analysis shows that sT and sT^D44N^ slightly reduced LT expression in U2OS cells. Replication efficiency data is shown in Fig. 1C. (**B**) Western blot analysis shows that both sT and sT^D44N^ enhanced LT expression in 293 cells. This is due to sT interaction with E3 ligase to reduce LT degradation in 293 cells. (**C**) qPCR analysis f. or the viral origin containing plasmid indicated that LT+ sT^D44N^ reduced replication by 47% compared to LT +sT, whereas LT^D44N^ +sT/sT^D44N^ reduces replication by 98% compared to LT+sT. This indicates that sT DnaJ domain contributes partially to sT enhancement in replication, and LT DnaJ domain contributes majorly to sT enhancement in replication.

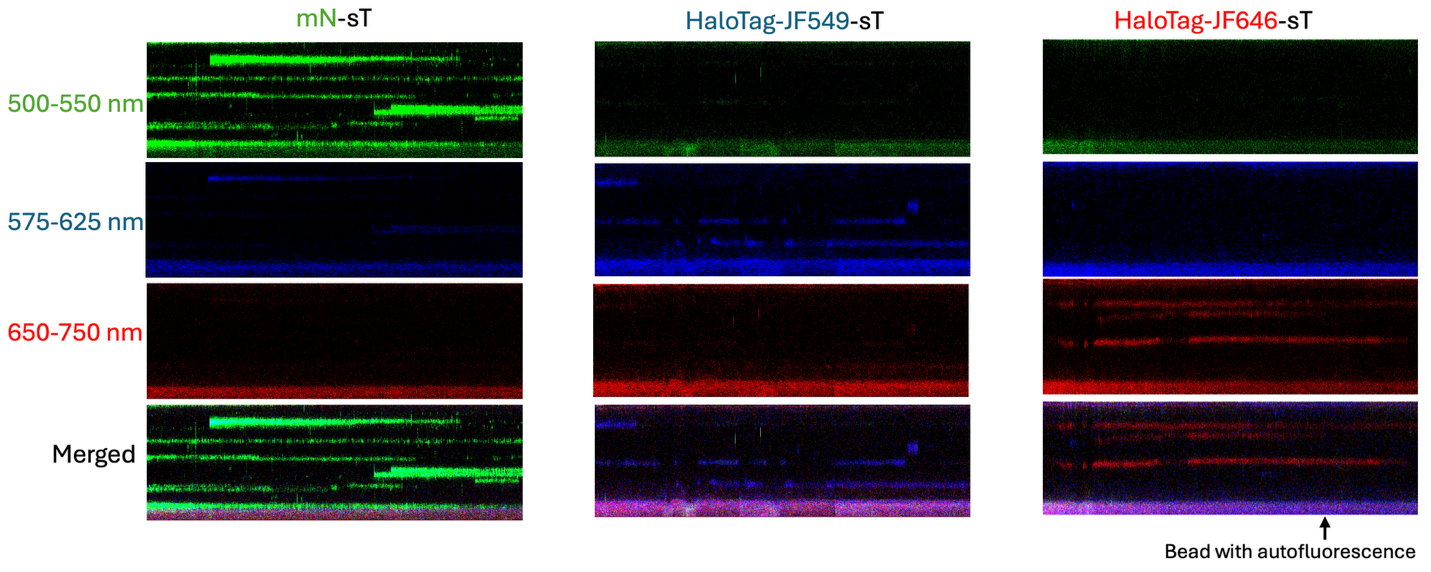

**Fig. S8. Spectral bleed-through control.** Representative kymographs of sT labeled with mNeongreen (left column), HaloTag-JF549 (middle column), or HaloTag-JF646 (right column). For all three conditions, the 488, 561, and 638 nm excitation lasers were simultaneously activated, and fluorescence was collected through three separate bandpass filters: 500–550 nm (mNeongreen), 575–625 nm (JF549), and 650–750 nm (JF646). Each column represents a single acquisition, displayed as a merged image (top row), followed by the individual 500–550 nm (second row), 575–625 nm (third row), and 650–750 nm (bottom row) detection channels. Polystyrene bead autofluorescence at the lower edge of each kymograph was visible in all three channels, serving as an internal reference for consistent image display and contrast settings. Images were displayed as described in Methods.

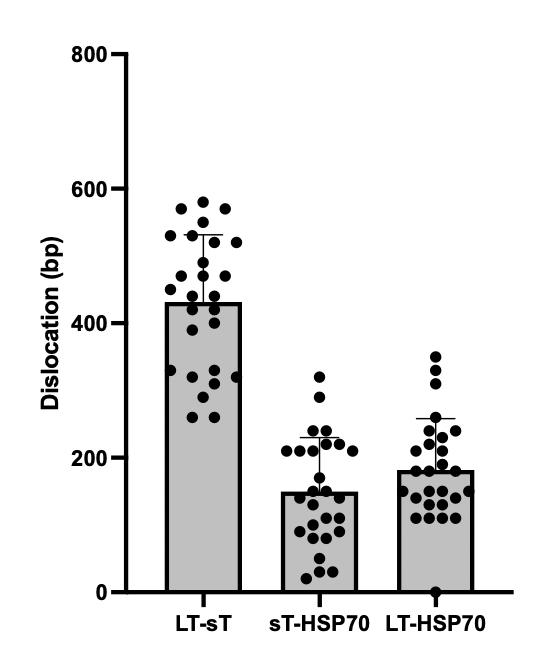

**Fig. S9. Spatial organization of the LT–sT–HSP70 complex.** Quantification of the distances between LT–sT, sT–HSP70, and LT–HSP70 in complexes formed with HaloTag-JF549-LT, HaloTag-JF646-sT, and EGFP-HSP70.

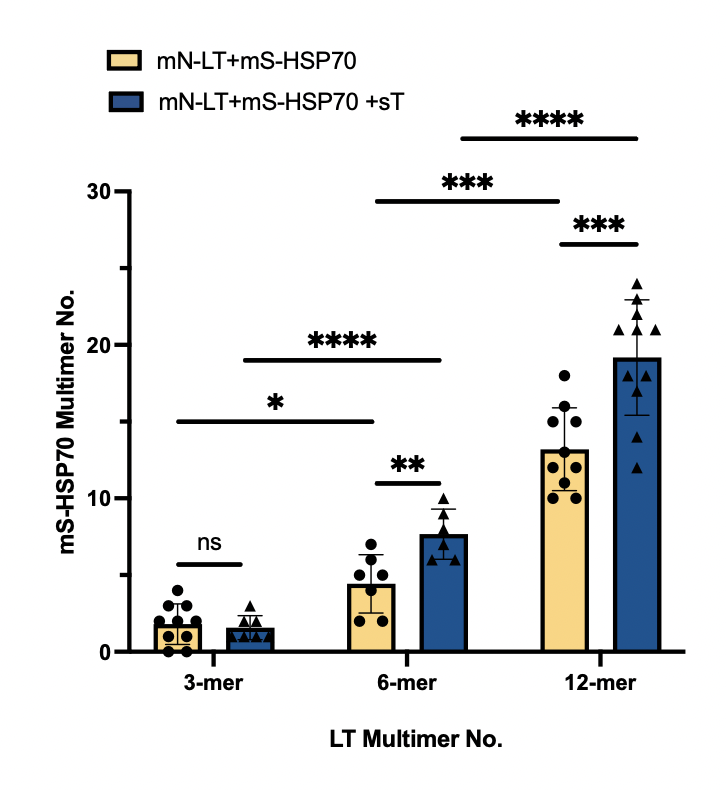

**Fig. S10. sT enhances HSP70 binding to the LT–sT complex.** Quantification of HSP70 molecules associated with mN-LT trimers, hexamers, and dodecamers, with and without sT. The number of HSP70 molecules per complex increased with LT multimerization level, with dodecameric complexes showing a median of approximately 20 HSP70 monomers. Addition of sT further enhanced HSP70 recruitment, consistent with the LT–HSP70–HSP70–sT bridging model.

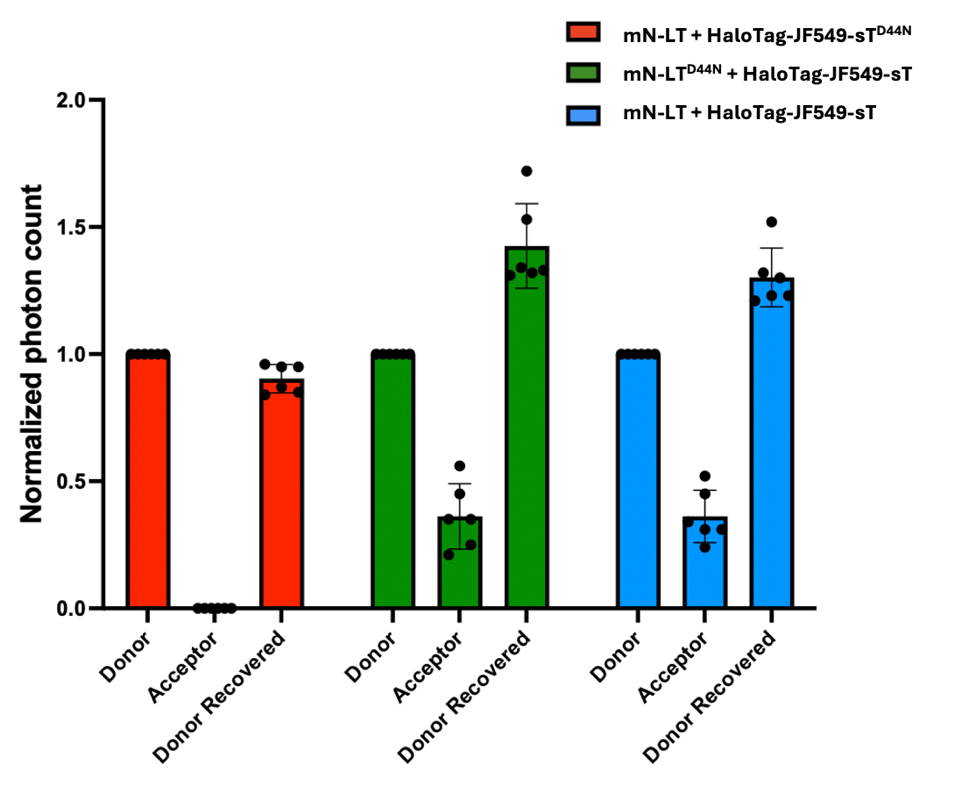

**Fig. S11. Normalized photon counts in single-molecule LT–sT FRET measurements.** Single-molecule Förster resonance energy transfer (FRET) was measured using mN-LT as the donor and HaloTag-JF549-sT as the acceptor. Photon counts were quantified for the donor, acceptor, and dequenched donor following acceptor photobleaching. Donor and acceptor photon counts were normalized to the initial donor fluorescence intensity. Six individual molecules were analyzed for each condition. Data are presented as mean ± SD. N=6 molecules per condition.

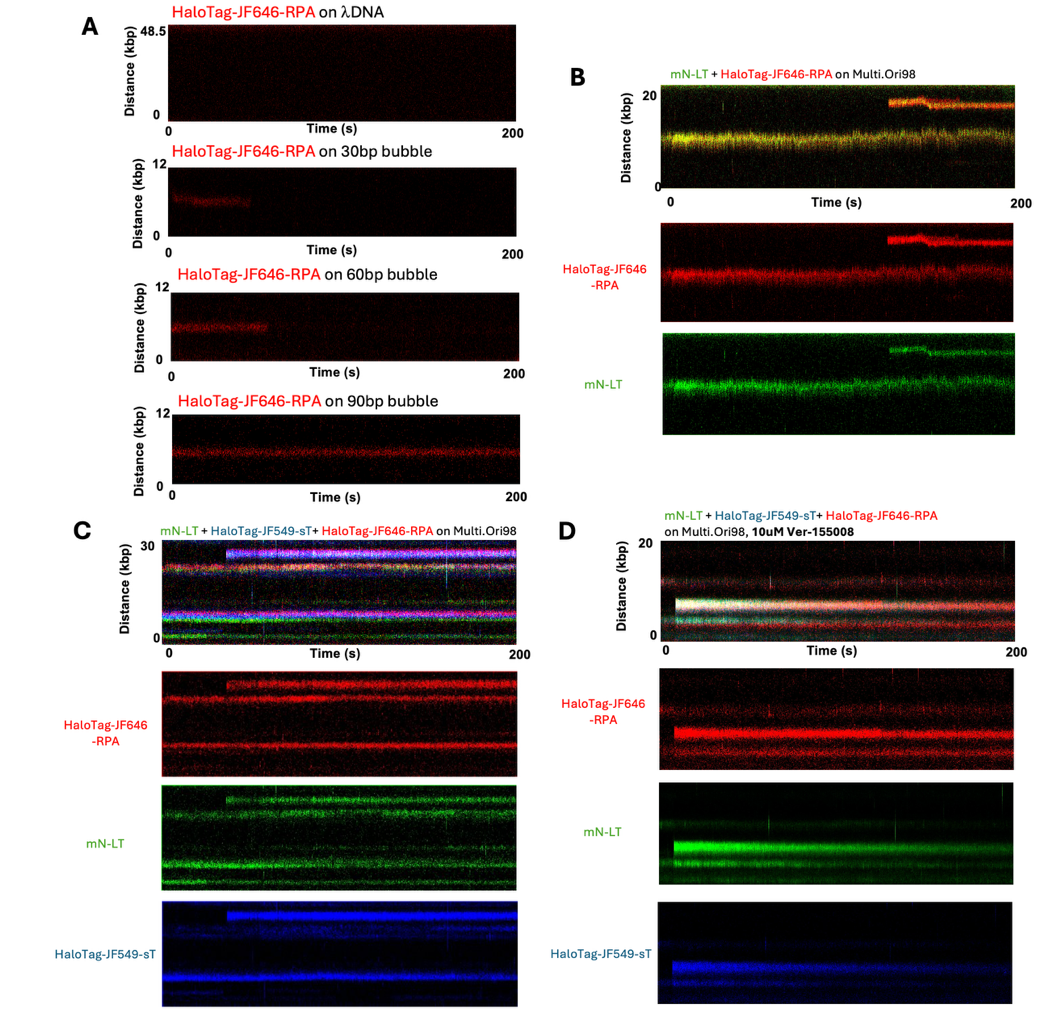

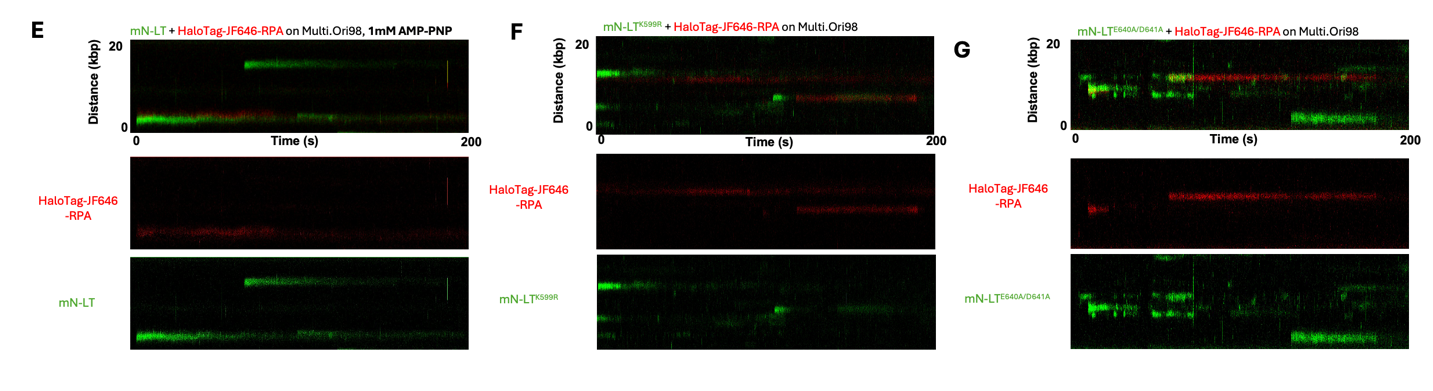

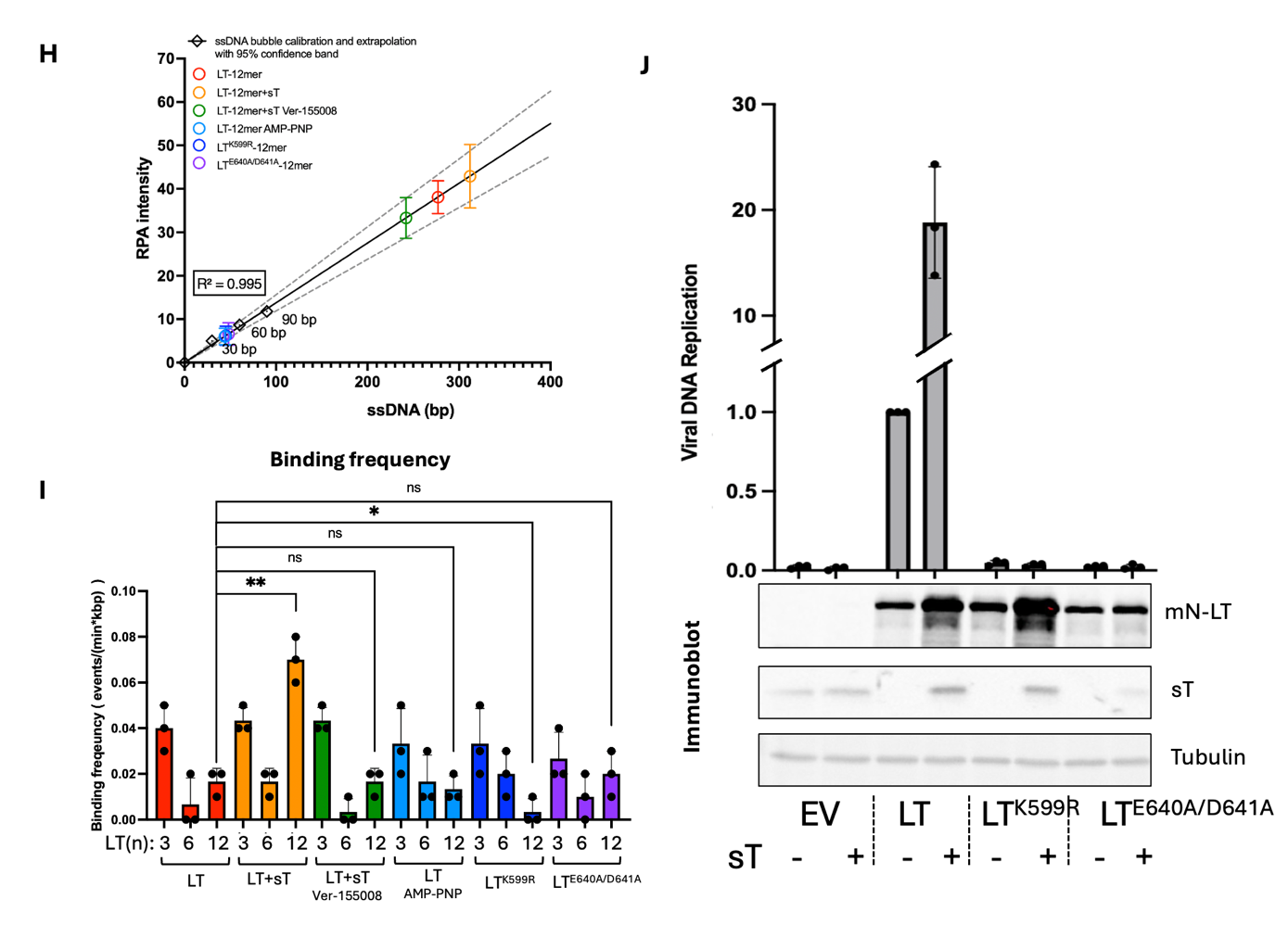

**Figure S12. RPA fluorescence-based analysis of DNA melting by MCV large T antigen.**

**(A)** Representative fluorescence intensity profiles of HaloTag-JF646-RPA (red) bound to DNA substrates containing 30-, 60-, or 90-bp poly(dA) bubble regions, serving as calibration standards for exposed ssDNA. λDNA serves as a negative control. **(B–G)** Representative fluorescence profiles of mN-LT (green) and HaloTag-JF646-RPA (red) bound to DNA (B); with HaloTag-JF549-sT (blue) (C); with HaloTag-JF549-sT and 10 μM Ver-155008 (D); with 1 mM AMP-PNP substituting ATP (E); with the Walker A mutant mN-LT^K599R^ (F); or with the Walker B mutant mN-LT^E640A/D641A^ (G). **(H)** Frequency of LT trimer, hexamer, and dodecamer assemblies across the conditions shown in (B–G). **(I)** Replicon assay demonstrating that Walker A (K599R) and Walker B (E640A/D641A) mutations abolish LT-mediated DNA replication, both in the absence and presence of sT co-expression. Immunoblot analysis confirms the expression of mN-LT and sT. Data are presented as mean ± SD from three independent replicates.

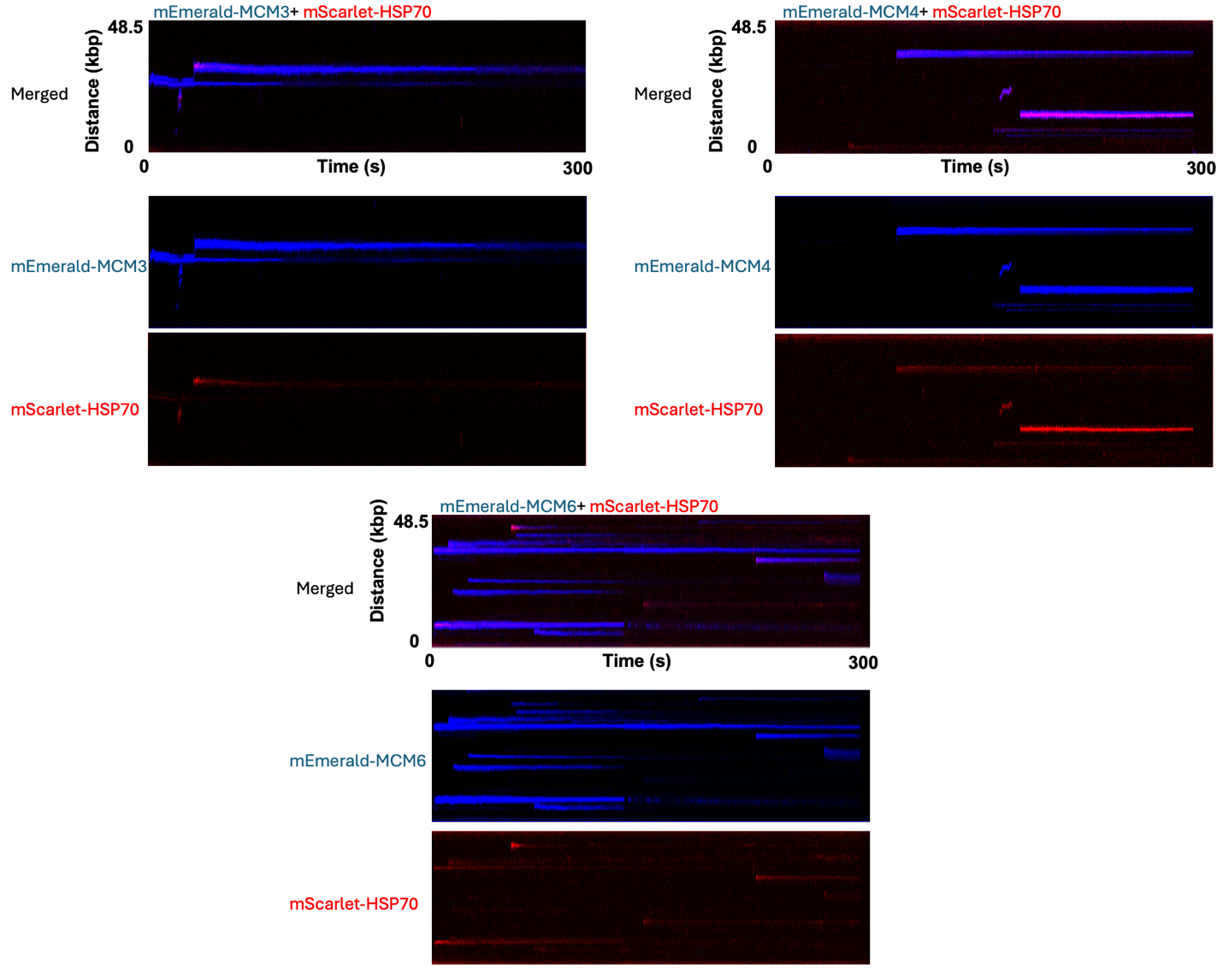

**Fig. S13. Individual fluorescence channels of MCM3, MCM4, and MCM6 colocalization with HSP70 on λ DNA.** Representative kymographs corresponding to the MCM3, MCM4, and MCM6 conditions shown in Fig. 5A. Each column shows the original merged kymograph and the corresponding individual mEmerald–MCM and mScarlet–HSP70 fluorescence channels from the same acquisition, illustrating colocalization of HSP70 with MCM3, MCM4, and MCM6 on DNA.

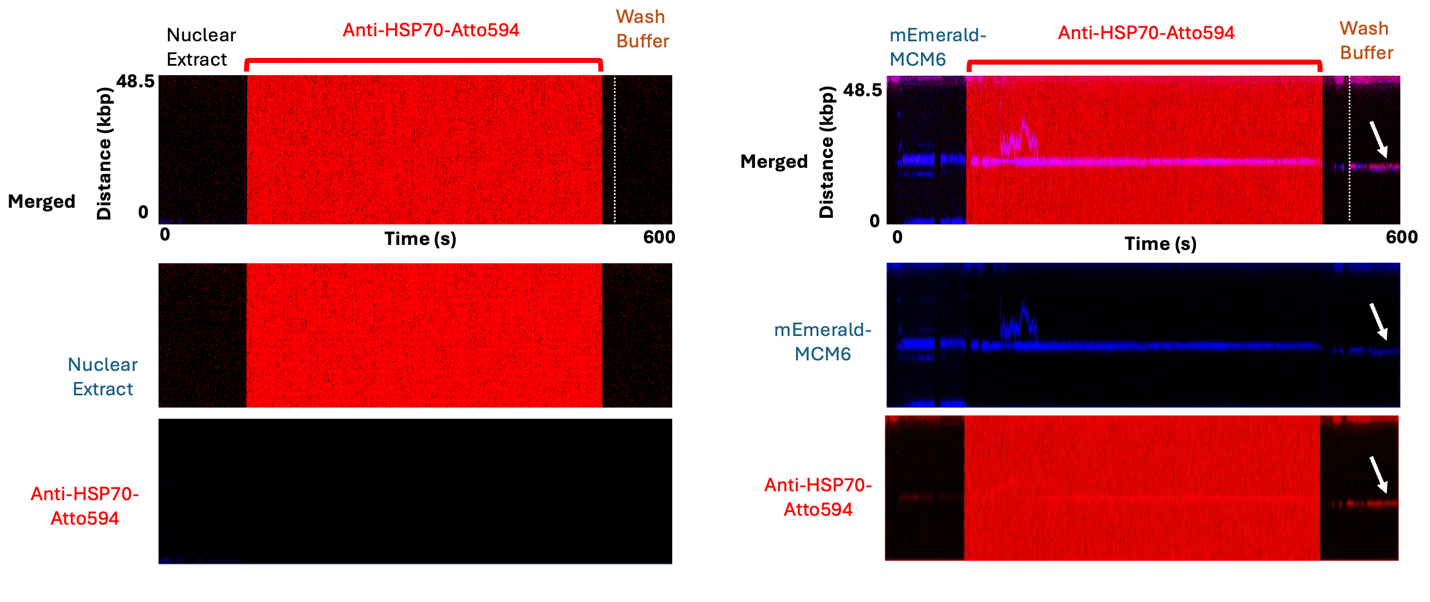

**Fig. S14. Individual fluorescence channels of MCM6 and endogenous HSP70 binding to λ DNA.** Representative kymographs showing mEmerald–MCM6 (blue) and endogenous HSP70 detected by anti-HSP70–Atto594 (red) on λ DNA using nuclear extracts with or without mEmerald–MCM6. Each panel shows the original merged kymograph and the corresponding individual fluorescence channels. Endogenous HSP70 colocalized with DNA-bound MCM6, whereas no detectable HSP70 binding was observed in the absence of MCM6. Both MCM6 and HSP70 signals persisted and remained colocalized following transfer to antibody-free wash buffer.

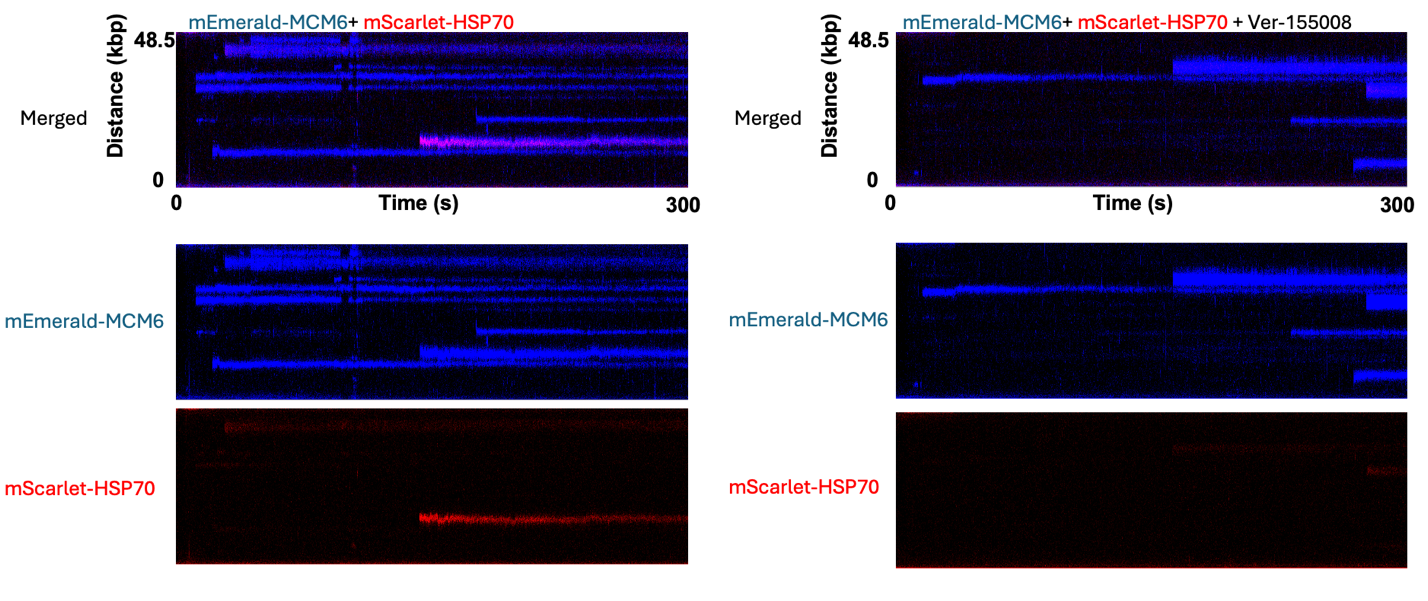

**Fig. S15. Individual fluorescence channels of MCM6 and HSP70 binding to λ DNA in the presence or absence of HSP70 inhibition.** Representative kymographs of mEmerald–MCM6 (blue) and mScarlet–HSP70 (red) binding to λ DNA under control conditions (left) or following pretreatment of nuclear extracts with 5 µM VER-155008 (right). Each panel shows the original merged kymograph and the corresponding individual mEmerald–MCM6 and mScarlet–HSP70 fluorescence channels from the same acquisition. HSP70 colocalized with DNA-bound MCM6 under control conditions, whereas VER-155008 treatment abolished detectable HSP70 recruitment to MCM6-bound DNA.

**Table S1. List and description of plasmid constructs.**

| **Construct** | **Function** | **Parental Vector** | **Chang-Moore Plasmid #** | **Chang-Moore Plasmid Name** |
| --- | --- | --- | --- | --- |
| pcDNA6 | Control Vector | pcDNA6.V5-HisB | 2892 | pcDNA6 modified |
| pcDNA6-LT | Expresses MCV codon-optimized LT in mammalian cells | pcDNA6 | 2956 | pcDNA6.MCV LTco |
| pcDNA6-LT(D44N) | Expresses MCV codon-optimized LT DNAJ domain mutation that eliminates HSC70 interaction | pcDNA6 | 3214 | pcDNA.LTco D44N |
| pcDNA6-sT | Expresses MCV codon-optimized sT in mammalian cells | pcDNA6 | 2915 | pcDNA6.MCV sTco |
| pcDNA6-sT(D44N) | Expresses MCV codon-optimized sT DNAJ domain mutation that eliminates HSC70 interaction | pcDNA6 | 2916 | pcDNA.sTco D44N |
| pmNeonGreen-C1 | Expresses mNeonGreen in mammalian cells | pmNeonGreen-C1 | 4732 | pmNeonGreen-C1 |
| pmNeonGreen-N1 | Expresses mNeonGreen in mammalian cells | pmNeonGreen-N1 | 4733 | pmNeonGreen-N1 |
| mNeonGreen-LT | Expresses mNeonGreen-fused codon-optimized LT in mammalian cells | pmNeonGreen-C1 | 4879 | mNeonGreen-MCV LTco |
| mNeonGreen-LT (D44N) | mNeonGreen-fused LT DNAJ domain mutation that eliminates HSC70 interaction | pmNeonGreen-C1 | 5324 | mN-LTco(D44N) |
| pmScarlet-C1 | Expresses mScarlet in mammalian cells | pmScarlet-C1 | 4738 | pmScarlet-C1 |
| LT-mScarlet | Expresses mScarlet-fused codon-optimized LT in mammalian cells | pmScarlet-C1 | 4780 | MCV LTco-mScarlet |
| pCR-Ori98 | MCV replication origin | pCR2.1-TOPO | 2285 | pCR.Ori339(98) |
| pMC-MCV-98bp | Amplifies Ori98 sites in pMC.BESPX backbone in bacteria for self-ligation and biotinylation to produce dsDNA tethers | pMC.BESPX | 4883 | pMC.BESPX-MCV-98bp |
| pEGFP-N1 | Expresses EGFP in mammalian cells | pEGFP-N1 | 2437 | pEGFP-N1 |
| EGFP-sT | Expresses EGFP-fused codon-optimized sT in mammalian cells | pEGFP-N1 | 4930 | sTco-GFP |
| EGFP-sT(D44N) | EGFP-fused sT DNAJ domain mutation that eliminates HSC70 interaction | pEGFP-N1 | 4934 | sTco.D44N-GFP |
| mNeonGreen-sT | Expresses mNeonGreen-fused codon-optimized sT in mammalian cells | pmNeonGreen-C1 | 4963 | mNeonGreen-MCV sTco |
| mNeonGreen-sT (D44N) | mNeonGreen-fused sT DNAJ domain mutation that eliminates HSC70 interaction | pmNeonGreen-C1 | 4964 | mNeonGreen-MCV sTcoD44N |
| mNeonGreen-sT (L142A) | mNeonGreen-fused sT mutation to inhibit PP2A interaction | pmNeonGreen-C1 | 4965 | mNeonGreen-MCV sTcoL142A |
| mNeonGreen-sT (A91-95) | mNeonGreen-fused sT LT stabilization domain mutation (substition with alanines) | pmNeonGreen-C1 | 4966 | mNeonGreen-MCV sTcoA91-95 LSD |
| mNeonGreen- sT (A83-88) | mNeonGreen-fused sT EP400-complex binding deficient mutation | pmNeonGreen-C1 | 4967 | mNeonGreen-MCV sTco83-88A |
| HaloTag-  LT | Expresses MCV LT fused with Halo tag in mammalian cells | pHTN-vector-stock_pLann | 5222 | Halo-MCV.LTco |
| HaloTag-  sT | Expresses MCV sT fused with Halo tag in mammalian cells | pHTN-vector-stock_pLann | 5223 | Halo-MCV.sTco |
| HaloTag-sT(D44N) | Expresses MCV sT fused with Halo tag DNAJ domain mutation that eliminates HSC70 interaction | pHTN-vector-stock_pLann | 5227 | Halo-MCV sTco(D44N) |
| mScarlet-HSP70 | Expresses mScarlet-fused HSP70 in mammalian cells | pmScarlet-C1 | 5261 | pmScarlet-HSP70(A1A) |
| EGFP-HSP70 | Expresses EGFP-fused HSP70 in mammalian cells | pEGFP-N1 | 5260 | pEGFP-HSP70(A1A) |
| mEmerald-MCM3 | Expresses mEmerald-fused MCM3 in mammalian cells | N/A | 5348 | mEmerald-MCM3(Mouse) |
| mEmerald-MCM4 | Expresses mEmerald-fused MCM4 in mammalian cells | N/A | 5349 | mEmerald-MCM4(Mouse) |
| mEmerald-MCM6 | Expresses mEmerald-fused MCM6 in mammalian cells | N/A | 5350 | mEmerald-MCM6(Mouse) |
| Halo-RPA | Expresses RPA14, Halo tagged RPA32, and RPA 70 in mammalian cells | N/A | 5502 | RPA70-IRES-Halo-RPA32-IRES-RPA14 |

**Table S2. Oligonucleotide sequences used to generate 30-, 60-, and 90-nt poly(A) DNA bubbles.**

| 30bpA-Bubble-S | /5Phos/CAACGTCATGCATGACGTAGGTTCGAACCTCGAATAAAAAAAAAAAAAAAAAAAAAAAAAAAAAAGAGTCAACTGTATCTAACGTCATGTACGTAG |
| --- | --- |
| 30bpA-Bubble-AS | /5Phos/ACCACTACGTACATGACGTTAGATACAGTTGACTCAAAAAAAAAAAAAAAAAAAAAAAAAAAAAAATTCGAGGTTCGAACCTACGTCATGCATGAC |
| 60bpA-Bubble-S | /5Phos/CAACGTCATGCATGACGTAGGTTCGAACCTCGAATAAAAAAAAAAAAAAAAAAAAAAAAAAAAAAAAAAAAAAAAAAAAAAAAAAAAAAAAAAAAGAGTCAACTGTATCTAACGTCATGTACGTAG |
| 60bpA-Bubble-AS | /5Phos/ACCACTACGTACATGACGTTAGATACAGTTGACTCAAAAAAAAAAAAAAAAAAAAAAAAAAAAAAAAAAAAAAAAAAAAAAAAAAAAAAAAAAAAATTCGAGGTTCGAACCTACGTCATGCATGAC |
| 90bpA-Bubble-S | /5Phos/CAACGTCATGCATGACGTAGGTTCGAACCTCGAATAAAAAAAAAAAAAAAAAAAAAAAAAAAAAAAAAAAAAAAAAAAAAAAAAAAAAAAAAAAAAAAAAAAAAAAAAAAAAAAAAAAAAAAAAAGAGTCAACTGTATCTAACGTCATGTACGTAG |
| 90bpA-Bubble-AS | /5Phos/ACCACTACGTACATGACGTTAGATACAGTTGACTCAAAAAAAAAAAAAAAAAAAAAAAAAAAAAAAAAAAAAAAAAAAAAAAAAAAAAAAAAAAAAAAAAAAAAAAAAAAAAAAAAAAAAAAAAAATTCGAGGTTCGAACCTACGTCATGCATGAC |
